# Use of photocaged molecular oxygen for time-resolved serial X-ray crystallography

**DOI:** 10.64898/2026.08.10.743912

**Authors:** Sofia Jaho, Arturo Landeros de la Isla, Danny Axford, Sinan Battah, James Beilsten-Edmands, Do-Heon Gu, Sam Horrell, Gabriel Karras, Marina Lučić, Aidan E. Meekings, Halina Mikolajek, Shigeki Owada, Hiroshi Sugimoto, Rachel Tang, Takehiko Tosha, Ivo Tews, Amy J. Thompson, Briony A. Yorke, Michael A. Hough, Jonathan A. R. Worrall, Robin L. Owen

## Abstract

Photocages offer an attractive means of synchronously triggering enzyme-substrate driven reactions in biological systems *in crystallo* expanding the reach of light-driven catalysis. We describe the application of photocaged molecular oxygen to trigger molecular oxygen binding in crystals of myoglobin under anaerobic conditions and follow structural changes using both serial synchrotron and serial femtosecond X-ray crystallography. This is enabled through use of fixed targets under anaerobic conditions, utilising thin polymeric films with low molecular oxygen permeability and validated by serially collecting deoxy myoglobin structures and complementary *in crystallo* UV-Vis spectroscopy. Release of molecular oxygen from the photocage and subsequent binding of the gaseous ligand is structurally visualised in oxygen-bound structures of myoglobin at 5 and 10 ms and various laser parameters. We present a robust workflow for enabling anaerobic room-temperature data collection of oxygen-sensitive samples on fixed targets and report the successful photo-release of caged molecular oxygen for time-resolved serial crystallography.

## Introduction

Time-resolved crystallography offers the promise of obtaining ensembles of structures, fully describing enzymatic pathways. The visualisation of short-lived intermediate states can reveal transient conformations or binding modes that may not be apparent from a single, static structure. Serial synchrotron crystallography (SSX) and serial femtosecond crystallography (SFX) offer attractive means of obtaining time-resolved data at synchrotrons and X-ray free electron lasers (XFELs) respectively and, in recent years, these approaches have increased in popularity (*1*). Time-resolved SSX and SFX remain experimentally challenging however, with stringent requirements on sample preparation, sample delivery and reaction initiation. Light is an attractive means of initiating reactivity offering high temporal resolution and negating the requirement to synchronously deliver substrate and allow it to diffuse uniformly through crystals. Only a small fraction (< 1%) of enzymes are naturally light activated however (*2*), and so approaches making use of photolabile amino acids (*3*, *4*) or photocages are required to make photoactivation more widely applicable.

While photocages have been used in crystallographic studies in the past to study slow reactions (*5–7*), their use in conjunction with serial crystallography allows faster turnover events to be probed. To date however, only a limited number of studies have utilised this combination of cage and experimental approach. A nitric oxide photocage has been used to access intermediate states of cytochrome P450nor (*8*), and initiate binding in a cytochrome c’ and a dye-decolourizing peroxidase (*9*). Mehrabi *et al*. utilised decaging of fluoroacetate in a time-resolved SSX experiment to reveal coupled allosteric motions between the two active sites of fluoroacetate dehalogenase (*10*). A further example is the use of photocaged zinc ions to initiate reactivity of a beta-lactamase (*11*), where laser light was used to release zinc ions and cleave the beta-lactam ring.

The decaging rate leading to compound release and the manner in which samples are serially delivered to the X-ray beam determine the time regimes that can be accessed. Tosha *et al*. utilised an LCP injector to obtain a 20 ms time point, with slower time points obtained via cryo SF-ROX (serial femtosecond rotation crystallography) and multi-crystal synchrotron diffraction (*8*). Fixed targets allow for an extremely broad range of timescales to be explored utilising a single sample delivery approach using either an excite and immediately collect or an excite and visit again strategy as demonstrated by both (*10*) (time span 30 ms – 30s) and (*9*) (100 µs - 1.4 s), or an excite and wait approach in combination with a mechanical chopper as utilised by (*11*) (20 ms – 4 s).

Wide uptake of photocaged molecular oxygen (O_2_) in structural biology has been restricted for two main reasons: ground-state O_2_ in a photocaged form is chemically unstable, while the necessity of preserving an anaerobic environment for both sample preparation and data collection is non-trivial. Experimental work on the synthesis of caged dioxygen carriers such as µ-superoxo or µ-peroxo-dicobalt(III) derivatives dates to the 1970s, with the µ-superoxo group reporting high quantum yield upon photolysis at pH values below the physiological range, while the µ-peroxo-dicobalt(III) group showed production of O_2_ in basic solutions upon UV irradiation (*12*). Building on this work (*13*) investigated several µ-peroxo-bridged complexes able to cage O_2_ at near physiological pH. Dioxygen was found to be released from a perchlorate salt of (µ-peroxo)(µ-hydroxo)bis[bis(bipyridyl)cobalt(III)] complex (HPBC) upon irradiation at 355 nm with a quantum yield of 0.04. Oxy-haemoglobin was formed from deoxy-haemoglobin upon photodissociation of the complex verifying that O_2_ was the primary photoproduct, providing new tools to study fast (ns or faster) biological reactions with spectroscopic probes. The same complex was used to study the oxidation of a fully reduced bovine *aa*_3_-type cytochrome *c* oxidase and the *ba*_3_ type (*14*), circumventing limitations of traditional stopped-flow experiments and spectroscopically resolving fast kinetic phases at µs – ms time resolution. Work by (*15*) reported a nitrate salt of HPBC with enhanced solubility compared to the perchlorate salt for time-resolved FT-IR spectroscopic studies of cytochrome *bo*_3_ oxidase from *Escherichia coli*. Photolysis at 308 nm resulted in a quantum efficiency of 0.4 and a stoichiometric release of dioxygen from HPBC.

Despite the considerable advances in the synthesis of caged dioxygen compounds and the widespread use in spectroscopic kinetic studies, one of the first documented attempts to study photolysis of the HPBC complex under cryogenic conditions was described by (*16*) paving the way for structural studies of cryo-trapped unstable intermediates of O_2_-dependent enzymes. The quantum yield at 100 K was determined to be 0.003 upon 266 nm laser flash, significantly lower compared to other reported values at room temperature. However, this low photolysis rate must be considered relative to the low catalysis rate at cryogenic temperatures where biological activity is effectively frozen out. More recently, (*17*) validated spectroscopically at room temperature the oxidation of reduced cytochrome *c* oxidase microcrystals from high (>3 mM per 355 nm single laser pulse) concentrations of O_2_ released by a caged compound demonstrating that the method could be extended to time-resolved X-ray crystallography, ensuring that at these high concentrations the reaction initiation is synchronous and homogeneous in the crystals.

Sample environments that maintain a room temperature anaerobic atmosphere are challenging to preserve. While in-vacuum or in-helium data collection is common for jet and extruder XFEL data collection (e.g. (*18*)), this is not often used at synchrotrons or with single crystal or serial fixed target approaches. Some early approaches mounted samples in capillaries sealed with either wax or epoxy resin (*19*). More recently, (*20*) enclosed fixed targets within multiple layers of thin film to enable anaerobic SSX of a hydroxylase, a DNA repair enzyme and an isopenicillin N synthase while (Bjelčić *et al.*, 2023) utilised a similar approach to study hemoglobin.

Myoglobin (Mb) is a small, globular heme protein capable of binding on the distal face of the heme gaseous ligands such as O_2_, CO and NO. It is therefore a model system well suited for characterisation of a O_2_-releasing photocage and associated experimental approaches and controls due to its structural simplicity, experimental accessibility, and defined ligand-binding chemistry. The ligand-binding is efficient and reversible, allowing kinetic processes such as ligand association, dissociation, and migration to be initiated and tracked with high precision ideal for time-resolved studies (*22*). The first successful time-resolved crystallography experiment on Mb revealed the structure of Mb photoproducts through the nanosecond timeframe during the process of heme and protein relaxation and ligand dissociation of CO at near-atomic resolution (*23*). This study demonstrated that structural changes following photolysis propagate over multiple timescales. Subsequent work extended the accessible temporal window from nanoseconds to milliseconds, identifying distinct intermediate states along the reaction coordinate. The advent of XFELs has extended the time resolution into the femtosecond regime, allowing investigation of the earliest electronic and nuclear events following ligand photolysis.

We describe a robust and reliable method for anaerobic data collection and reaction initiation using fixed targets at both synchrotron and XFEL sources enabling straightforward systematic structural studies of oxygen-sensitive enzymes. The proposed workflow utilises polymeric films with low-oxygen permeability ensuring anaerobic conditions are maintained for the duration of the experiment. Horse heart myoglobin (hhMb) was used as a model system to validate the robustness and reproducibility of the approach with *in crystallo* UV-Vis spectroscopy tracking the oxidation state of the heme iron in reduced and oxidised microcrystals used as additional validation. Following extensive iterative optimisation of sample preparation, we report the first anaerobic SSX and SFX structures of deoxy hhMb at room temperature showcasing the reliability of the anaerobic data collection. This anaerobic sample delivery approach paves the way for application of a O_2_ releasing photocage for studying structural dynamics with serial crystallography. We successfully used different illumination conditions to photo-release caged O_2_ and track the subsequent binding in hhMb with SSX and SFX on the millisecond timescale. Finally, we describe the implementation of a series of control experiments, including a checkerboard data collection scheme, emphasizing the importance of such tools for cross-validation of the anaerobicity of our method, time resolution of the structural snapshots and successful photocage activation.

## Methods

### Synthesis of photocaged molecular oxygen

The (µ-peroxo)(µ-hydroxo)bis[bis(bipyridyl)cobalt(III)] (HPBC) perchlorate salt complex was synthesized following the procedure described in (*15*) with minor alterations. For the synthesis, 2 g (13 mmol) of 2,2′-bipyridine dissolved in 20 mL of ethanol was added to 1.9 g (6.4 mmol) of cobalt(II) nitrate hexahydrate (Co(NO_3_)2·6H_2_O) dissolved in 20 mL of water. The pH of the mixture was adjusted to 9.2 using 1 M NaOH, and all subsequent steps were performed in the dark by wrapping the reaction flask with aluminium foil. The mixture was stirred while oxygen gas was bubbled through for 20 minutes; the pH was then readjusted to 9.6, and oxygenation continued for further 20 minutes. Crystallization was induced by adding 2 mL of a 6 M sodium perchlorate solution (in 50 % aqueous ethanol) and incubating the mixture at 4 °C overnight. The resulting black crystals were collected by suction filtration, washed with chilled ethanol, and dried under vacuum for 6 hours, yielding 70 % of the dried salt. The spectroscopic properties of the final synthesised compound matched those reported in the literature (*15*) with wavelength maxima (λ_max_) measured at 460 nm, 395 nm, 314 nm, 304 nm (under dark conditions) and 293 nm (after white light illumination). UV-Vis spectra of the HPBC cage prior to, and following, exposure to white light are shown in Figure S1a. Figure S1b shows spectral changes of the cage mixed with deoxy-hhMB upon exposure to white light.

### Protein preparation and crystallisation

Horse heart myoglobin (hhMb) solution and microcrystals were prepared under anaerobic conditions in a glovebox (Belle Technology) maintaining O_2_ concentrations of < 10 ppm. The lyophilised protein powder (Sigma-Aldrich) and all degassed solutions were ported into the glovebox ≥ 24 h in advance, and 150 – 200 mg of protein powder were dissolved in 2.5 mL of 50 mM Tris/HCl pH 7.6 and 1.9 M ammonium sulphate (solution A). Solid sodium dithionite was added in excess until the colour of the protein solution turned from brown to red (spectra shown in Figure S2), indicating successful reduction of the heme iron to give the deoxygenated form. To remove excess of dithionite from the protein solution, a PD10 desalting column (Cytiva) was used. After washing the column with solution A, the hhMb solution was applied followed by the addition of 3.5 mL of solution A to elute fractions of hhMb that were collected at a protein concentration of 8 – 10 mg mL^-1^. The reduced hhMb aliquots were concentrated to a final concentration of approximately 40 mg mL^-1^ for subsequent crystallisation. Protein concentrations lower than 40 mg mL^-1^ were found not to yield any crystals, while concentrations higher than 60 mg mL^-1^ induced protein precipitation.

hhMb crystals were grown in batch by mixing the concentrated protein solution (40 mg mL^-1^) with saturated ammonium sulphate solution (4 M) in a 1:1 ratio. 5-10 µL of seed stock was then added. The seed stock solution was prepared by addition of solid ammonium sulphate to the concentrated protein solution until visible precipitation was observed (*24*). A mixture of thin plates and needle-shaped crystals (Figure S3) grew within 1 – 4 weeks depending on the final protein concentration, with average crystal sizes of 30 × 30 × 10 or 50 × 10 × 10 µm^3^, respectively. Prior to X-ray diffraction experiments, microcrystals were transported to a glovebox (Coy Laboratory Products) for chip loading and subsequent data collection. Excess sodium dithionite was added to the microcrystals before each experiment to minimise potential oxidation due to accidental exposure to atmospheric oxygen. The quality of diffraction from crystals was found to improve over time with crystals diffracting more reliably and to higher resolution after being left in the glovebox under anaerobic conditions for some time (4-5 months) after crystallisation.

### Anaerobic setup

An approach developed from that described by (*20*) was used to prepare anaerobic fixed targets. Rather than the multiple film layer approach described by Rabe, a single layer of 12.5 µm EVAL^TM^ EVOH EF-F film (Kuraray; https://eval.kuraray.com/en-emea/products/eval-monolayer-film/) was used to seal each side of all fixed targets. As previously described, EVAL allows high transmission of 308 nm laser light while maintaining a humid crystal environment and minimizing X-ray scatter (*9*). EVAL also provides an extremely effective barrier to O_2_ making it suitable for anaerobic experiments, and simplifies the approach described by Rabe *et al*. as a single layer of film is sufficient to maintain the anaerobicity of the fixed targets. EVAL was found to be better suited to fixed target experiments than other possible sealing films, such as polyvinylidene chloride (PVDC), due to its low X-ray scatter over the majority (∼10-1.8 Å) of the resolution range of interest, and high optical transmission at 308 nm, the wavelength used for decaging. EVAL film ensures that the sample remains anaerobic for the whole duration of data collection which is approximately 10 min at beamline I24 (Diamond Light Source, UK) and 15 min at the SACLA XFEL (Japan).

In order to confirm that an anaerobic sample environment could be maintained from sample preparation, transfer and throughout serial X-ray experiments, hhMb was reduced in the presence of excess sodium dithionite (see Protein preparation and crystallisation). Characteristic UV-Vis spectra of reduced and oxidised hhMb crystals on EVAL films under excess dithionite are shown in Figures 1b and S4. hhMb could also be successfully reduced using only 50 mM of sodium dithionite as shown in Figure S5b. Both SSX and SFX experiments were carried out in the presence of excess dithionite however in order to eliminate the possibility of partially reduced sample. This was key when working with the O_2_ photocage to ensure fully reduced conditions for maximal oxygen binding upon decaging.

**Figure 1.**
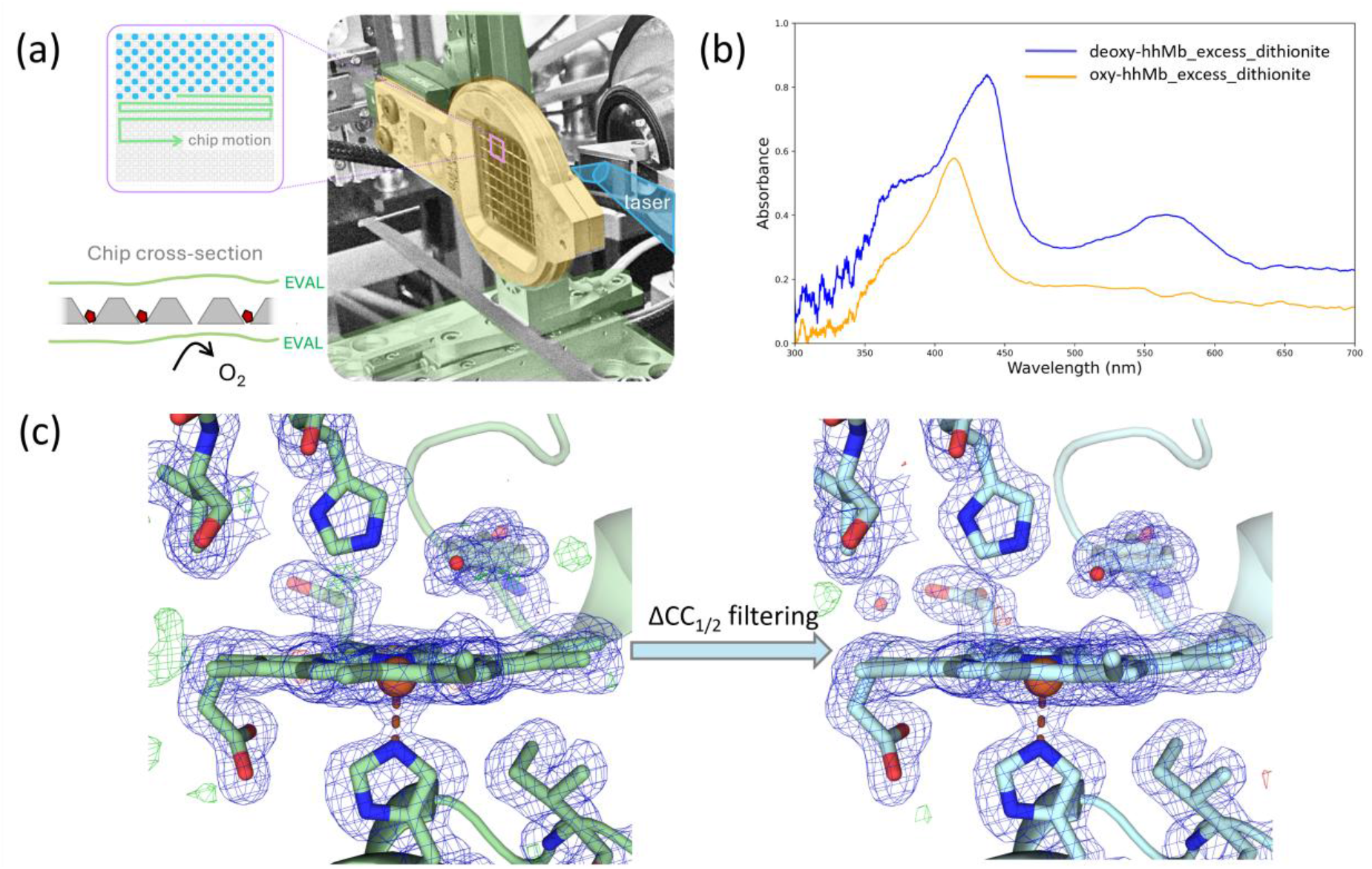
Schematic of setup (a) illustrating data collection hardware at I24, checkerboard data collection – highlighted by alternating blue and white apertures - and mode of anaerobic sealing. (b) UV-Vis spectra obtained from hhMb microcrystal slurries of reduced deoxy-(Fe^2+^) and oxy-(Fe^3+^) hhMb microcrystals in the presence of excess sodium dithionite (orange and blue lines, respectively) mounted in ‘chipless chips’ of EVAL film. (c) Impact of ΔCC_1/2_ filtering on SSX electron density obtained from deoxy hhMb. Processing and refinement statistics are given in Table 2. 2Fo-Fc and Fo-Fc maps contoured at 1 and ±3 σ, respectively.

Fixed targets (“chips”) were loaded in an anaerobic chamber (Coy Laboratory Products) with maximum oxygen concentration of < 10 ppm. Chips, EVAL film and holders were ported in the chamber and allowed to degas for several hours (typically a minimum of 2-3 h or ideally overnight) before the experiment. On average the batches of hhmb microcrystals had a density of 10^7^ – 10^8^ crystals mL^-1^. At these high crystal density values, 50 µL of crystal slurry was found to be sufficient for chip loading ensuring a good hit rate as reported in (*25*).

Sample preparation, chip loading and data collection for experiments using the O_2_ photocage were performed under dim red lighting to avoid accidental or premature decaging due to ambient light. 10 mg of photocage, enveloped in aluminium foil, were ported in the glovebox prior to the experiment and mixed with 1 mL of the crystal buffer (1: 1 mixture of solution A and 4 M ammonium sulphate). hhMb microcrystals were left to settle, and the supernatant was removed and replaced with the same volume of the cage solution and left to soak for 10 minutes. Small volumes of crystals soaked in cage solution sufficient for loading 1-2 chips were freshly prepared prior to loading each chip to minimise premature decaging and O_2_ release due to the reducing agent (sodium dithionite) or any other buffer or precipitant. A DS-11 (DeNovix) spectrometer was used inside the glovebox to measure spectra of hhMb microcrystals at various steps during the sample preparation ensuring that the sample was reduced when mixed with dithionite and that it stayed in the reduced form when mixed with the photocage prior to any photoexcitation experiment. Once the chips were loaded, they were placed on the holders with EVAL film, ported out of the glovebox and transferred to the beamline in a light tight box.

Serial crystallography experiments were performed both at beamline I24, Diamond Light Source and at BL2 EH3, SACLA using a fixed target approach and the same silicon chips and near identical hardware. In brief, chips were aligned to the co-incident X-ray and laser beams using an on-axis microscope and a co-ordinate system detailing the position and angular orientation of each chip with respect to the beamline created *via* a graphical user interface (*26*). This co-ordinate system was uploaded to a DeltaTau Geobrick, which controlled fixed target motion, and chips were then rastered through the X-ray beam at ∼40 Hz (I24) or matching the repetition rate of SACLA (30 Hz) pausing at each aperture for X-ray exposure.

In both substrate and light driven time-resolved experiments, the generation of control data is essential to confirm the validity of timepoints and lack of cross contamination (*27*, *28*). This is often achieved in both jet/extruder and fixed target serial data collection through use of alternating dark and laser-pumped exposures. We here minimise light leakage and unintended photocage release through use of opaque silicon chips with apertures deeper than the crystal size and spacing greater than the laser focal spot, and use of checkerboard photoexcitation. In this data collection mode, laser illumination is triggered only for every other aperture of the fixed targets leading to two distinct datasets from each single chip, a control, or dark, dataset and a light-activated dataset (Figure 1a). Checkerboard excitation reduces the rate of generation of light-driven data by a factor of two but acts as an extremely valuable in-experiment control as the scheme allows for rapid assessment of cross-contamination between adjacent wells on a chip; in laser-activated experiments, if the dark dataset is contaminated by bound O_2_, then the gaseous ligand has diffused to other wells, contaminating also the light dataset by an inaccurate time point. The efficiency of photoexcitation in all experiments was maximised by ensuring all chips were orientated such that the open/wider end of apertures faced the laser.

### I24 data collection

At I24 data were either collected at 12.4 keV and recorded using a Pilatus3 6M detector, or at 20 keV using an Eiger2 9M CdTe detector. Pilatus data collections used 20 % transmission and a 10 ms exposure time, while for Eiger data collections the exposure time was 5 ms and full beam was used. For all experiments, the X-ray beam was defocussed to 20 × 20 μm^2^. The incident fluxes at 12.4 and 20 keV were 1.0 × 10^12^ ph s^-1^ and 1.9 × 10^12^ ph s^-1^, and the estimated diffraction weighted dose (DWD) absorbed by each crystal was 2.8 kGy and 0.94 kGy at 12.4 and 20 keV respectively. Doses were calculated by RADDOSE-3D (*29*). Time-resolved experiments at I24 utilised the PORTO laser system setup as described by (*9*) using an excitation wavelength of 308 nm. The laser was focussed down to 60 μm diameter and samples illuminated at 5 kHz over 5 ms (i.e. 25 pulses total) with an incident energy of 0.47 µJ per pulse.

### SACLA data collection

At SACLA data were collected using an X-ray energy of 11 keV and a focal spot size of ∼1.4 × 1.4 μm^2^. X-ray pulses were attenuated by 64 % using a 200 μm aluminium filter. The average dose in the exposed region (ADER) for each crystal was 520 kGy. Dose calculated using RADDOSE-XFEL (*30*) and assumes an (attenuated) pulse energy of 110 μJ/per pulse at sample and a pulse duration of 10 fs. While this dose is some two orders of magnitude greater than the DWD delivered at I24, we note that the DWD calculated for synchrotron experiments and the time-resolved ADER for XFEL experiments are not directly comparable and the structures determined at SACLA are expected to be free from any damage artefacts (*31*). Crystals were optically excited using a single 5 ns laser pulse using a wavelength of 308 nm. The laser was focussed to ∼70 μm diameter and the laser energy used for each experiment is given in Table 1. Live hit-rates were monitored and on-the-fly processing performed using the Cheetah/CrystFEL pipeline (*32*). SACLA checkerboard experiments were separated based on photodiode readings at sample position. This was done using the SACLA provided DataAccessUserAPI to read the photodiode values for each image tag into python (3.7). Images were sorted into “dark” and “light” based on a threshold photodiode reading of 0.05 V. Using the h5py library, the sorted tags were matched to h5 image files. Separate .lst files were saved for “dark” and “light” (each line contained a h5 frame with a specific image tag) and used as the input to CrystFEL.

**Table 1.** Data processing and refinement statistics for O_2_-bound hhMb SFX structures obtained using an O_2_ photocage, a deoxy structure without cage, a control structure (deoxy-hhMb soaked in O_2_ cage and a probe-pump data collection scheme). The last two columns include the statistics for the checkboard collection scheme with the two datasets described as “checker dark” and “checker light”. Values in parentheses refer to the outermost resolution shell. For all data the space-group was P12_1_1. All SFX structures were indexed, integrated and scaled using CrystFEL 0.11.1 (*34*).

| Dataset Name | Deoxy SFX | Oxy-cage SFX<br>(10 ms) | Oxy-cage SFX<br>(10 ms) | Control SFX | Checker dark SFX<br>(100 μs) | Checker light SFX<br>(100 μs) |
| --- | --- | --- | --- | --- | --- | --- |
| X-ray source | SACLA | SACLA | SACLA | SACLA | SACLA | SACLA |
| Laser energy / μJ | n/a | 28.6 | 191 | 30 (pump after probe at 10 ms) | 35 | 35 |
| Processing | CrystFEL | CrystFEL | CrystFEL | CrystFEL | CrystFEL | CrystFEL |
| Resolution ($\text{\AA}$) | 34.54 - 1.70<br>(1.73 - 1.70) | 61.38 - 1.70<br>(1.73 - 1.70) | 61.27 - 1.70<br>(1.73 - 1.70) | 61.37 - 1.7<br>(1.73 - 1.70) | 26.70 - 1.70<br>(1.73 - 1.70) | 26.71 - 1.70<br>(1.73 - 1.70) |
| Unit cell ($\text{\AA}$) | 64.33 28.55 35.72<br>90 107.55 90 | 64.36 28.55 35.68<br>90 107.51 90 | 64.18 28.59 35.66<br>90 107.32 90 | 64.31 28.65 35.71<br>90 107.38 90 | 64.40 28.73 35.79<br>90 107.26 90 | 64.45 28.74 35.81<br>90 107.29 90 |
| Merged crystals | 10827 | 5825 | 3008 | 18211 | 26789 | 26410 |
| Total observations | 615660 (19086) | 377596 (11792) | 168527 (5058) | 628527 (20170) | 1101463 (37015) | 1096682 (36253) |
| Unique reflections | 13927 (658) | 13922 (660) | 13855 (652) | 13775 (655) | 13802 (685) | 13841 (679) |
| Completeness | 100 (100) | 100 (100) | 99.67 (99.24) | 98.58 (99.85) | 98.03 (100) | 98.18 (100) |
| Multiplicity | 44.2 (29.0) | 27.1 (17.9) | 12.1 (7.8) | 45.6 (30.8) | 79.8 (54.0) | 79.2 (53.4) |
| CC<sub>1/2</sub> | 0.942 (0.568) | 0.908 (0.638) | 0.830 (0.564) | 0.944 (0.350) | 0.960 (0.779) | 0.965 (0.849) |
| R<sub>split</sub> | 0.186 (0.408) | 0.234 (0.440) | 0.314 (0.621) | 0.174 (0.350) | 0.146 (0.267) | 0.142 (0.272) |
| I/$\sigma$(I) | 6.60 (3.40) | 5.59 (2.75) | 4.53 (2.26) | 6.58 (4.67) | 7.47 (5.41) | 7.68 (5.40) |
| R<sub>work</sub> | 0.181 (0.262) | 0.182 (0.282) | 0.208 (0.320) | 0.192 (0.296) | 0.191 (0.339) | 0.190 (0.311) |
| R<sub>free</sub> | 0.228 (0.220) | 0.230 (0.298) | 0.247 (0.332) | 0.246 (0.321) | 0.244 (0.343) | 0.239 (0.322) |
| RMSD bond length ($\text{\AA}$) | 0.0054 | 0.0140 | 0.0045 | 0.0135 | 0.0130 | 0.0104 |
| RMSD bond angles ($^{\circ}$) | 1.350 | 2.312 | 1.288 | 2.119 | 2.142 | 1.841 |
| Ramachandran favoured (%) | 94.08 | 96.05 | 96.05 | 94.74 | 95.39 | 95.39 |
| O<sub>2</sub> occupancy | n/a | 0.2 | 0.35 | n/a | n/a | n/a |
| PDB ID | pdb_000032YU | pdb_000032YV | n/a | pdb_000032ZN | pdb_000032ZP | pdb_000032ZO |

### Data analysis

Diffraction data were indexed, integrated and scaled using either xia2.ssx software version 3.29.0 (*33*) or the CrystFEL software suite version 0.11.1 (*34*). Tables 1 and 2 detail which package was used for which datasets. For the data processed with CrystFEL, peak finding was performed with Peakfinder8 (*35*), the *Xgandalf* algorithm (*36*) was used for indexing, and peak integration was performed using the *--int-radius* method and optimised for each beamtime. Detector geometry was refined using the *detector-shift* and the Millepede-II method (*37*). All integrated reflections were scaled and merged using *partialator* using the *xsphere* model (*38*).

**Table 2.** Data processing and refinement statistics for a deoxy-hhMb SSX structure, a control structure (deoxy-hhMb soaked in O_2_ cage and data collected without any pump scheme) and O_2_-bound hhMb SSX structures obtained using an O_2_ photocage and the checkboard collection scheme with the two datasets described as “checker dark” and “checker light”. Values in parentheses refer to the outermost resolution shell. For all data the space-group was P12_1_1. All data in Table 2 were collected using an X-ray energy of 20.0 keV. All SSX structures were indexed, integrated and scaled using xia2.ssx (*33*), apart from the control structure which was processed with CrystFEL 0.11.1 (*34*). Crystals were photo-excited using 25 laser pulses delivered over 5 ms.

| Dataset Name | Deoxy SSX - $\Delta CC_{1/2}$ filtered | Deoxy SSX - $\Delta CC_{1/2}$ unfiltered | Control SSX | Checker dark SSX (5 ms) | Checker light SSX (5 ms) |
| --- | --- | --- | --- | --- | --- |
| X-ray source | I24 | I24 | I24 | I24 | I24 |
| Laser energy per pulse (total) / μJ | n/a | n/a | n/a | 0.47 (11.7) | 0.47 (11.7) |
| Processing | xia2.ssx | xia2.ssx | CrystFEL | xia2.ssx | xia2.ssx |
| Resolution ($\text{\AA}$) | 61.82 - 1.80<br>(1.83 - 1.80) | 61.82 - 1.80<br>(1.83 - 1.80) | 61.34 - 2.00<br>(2.05 - 2.00) | 61.52 - 1.80<br>(1.83 - 1.80) | 61.53 - 1.80<br>(1.83 - 1.80) |
| Unit cell ($\text{\AA}$) | 64.70 28.82 35.97<br>90 107.17 90 | 64.70 28.82 35.97<br>90 107.17 90 | 64.44 28.77 35.77<br>90 107.06 90 | 64.37 28.78 35.78<br>90 107.12 90 | 64.37 28.78 35.78<br>90 107.11 90 |
| Merged crystals | 5638 | 5839 | 19731 | 8849 | 8961 |
| Total observations | 338803 (10645) | 367986 (11939) | 863112 (28206) | 641861 (21002) | 648115 (21435) |
| Unique reflections | 12027 (570) | 12029 (572) | 8403 (414) | 11899 (576) | 11899 (577) |
| Completeness | 100 (100) | 100 (100) | 96.23 (100) | 100 (100) | 100 (100) |
| Multiplicity | 28.2 (18.7) | 30.6 (20.9) | 102.7 (68.1) | 53.9 (36.5) | 54.5 (37.1) |
| $CC_{1/2}$ | 0.979 (0.449) | 0.807 (0.392) | 0.947 (0.350) | 0.984 (0.639) | 0.983 (0.644) |
| $R_{\text{split}}$ | 0.159 (0.813) | 0.316 (0.818) | 0.186 (0.806) | 0.121 (0.523) | 0.123 (0.530) |
| $I/\sigma(I)$ | 5.10 (1.00) | 5.10 (1.40) | 5.00 (0.79) | 6.50 (1.20) | 6.40 (1.30) |
| $R_{\text{work}}$ | 0.183 (0.324) | 0.214 (0.296) | 0.198 (0.456) | 0.170 (0.264) | 0.167 (0.276) |
| $R_{\text{free}}$ | 0.242 (0.388) | 0.270 (0.386) | 0.244 (0.379) | 0.230 (0.278) | 0.232 (0.301) |
| RMSD bond length ($\text{\AA}$) | 0.0135 | 0.0135 | 0.0052 | 0.0133 | 0.0131 |
| RMSD bond angles ($^{\circ}$) | 2.079 | 2.233 | 1.338 | 2.199 | 2.111 |
| Ramachandran favoured (%) | 94.08 | 95.39 | 94.74 | 94.74 | 94.08 |
| O<sub>2</sub> occupancy | n/a | n/a | n/a | 0.45 | 0.85 |
| PDB ID | pdb_000032ZM | n/a | pdb_000032ZD | pdb_000032ZW | pdb_000033AB |

For the data processed with xia2.ssx, a detector geometry for each condition was refined using the default spotfinding and indexing algorithms. For the complete processing, hit rates and indexing rates were increased by using a spotfinding dispersion gain of 0.85 and by attempting all available indexing methods (fft1d, ffbidx, pink_indexer, low_res_spot_match and real_space_grid_search). Checkerboard data were split based on even/odd image number after integration and before scaling. Integrated reflections were initially scaled and merged with xia2.ssx_reduce, followed by 6 cycles of ΔCC_1/2_ filtering with dials.scale, using a filtering level of 3 standard deviations (stdcutoff=3.0) (*39*). ΔCC_1/2_ filtering removed between 2.1 % and 3.4 % of crystals in total from the merged datasets.

Phases were obtained with molecular replacement using PHASER (*40*) or MOLREP (*41*) with PDB entry 5D5R (*42*) as a search model. Models were iteratively refined in CCP4i2 (*43*) using REFMAC (*44*), Phenix (*45*) and Coot (*46*) for model building. Scaling and refinement statistics are summarised in Tables 1 and 2. Difference maps and Polder maps were calculated using Phenix (*47*). Validation was performed with MolProbity (*48*), PDB-REDO (*49*) and OneDep wwPDB (*50*) and *pdb_extract* (*51*) was used to prepare files for deposition in the Protein Data Bank. Structural figures were generated with PyMOL (Schrödinger).

## Results

### Deoxy-myoglobin SSX and SFX structures

Room temperature anaerobic deoxy hhMb structures obtained by both SSX (1.8 Å resolution) and SFX (1.7 Å resolution) are shown in Figures 2a and 2b, respectively. To our knowledge these represent the first such room temperature anaerobic structures obtained from Mb. To obtain these structures, it was necessary to maintain strict anaerobic conditions throughout sample preparation, chip loading and data collection. A consistently anaerobic environment is essential to preserve the reduced deoxy state of the heme iron, which in turn is key for subsequent experiments deploying the O_2_ photocage. Our initial experiments highlighted the importance of developing a thorough and reliable workflow for sample handling and delivery ensuring anaerobicity when working with oxygen-sensitive enzymes. Re-oxidation of the hhMb iron through partial exposure to ambient conditions in early experiments emphasized this necessity. Figure S6 shows the O_2_-bound SSX and SFX structures of hhMb, both at 1.7 Å resolution, where crystals were either not fully reduced or accidentally exposed to the atmosphere. The heme site evidentially shows O_2_ binding with occupancies estimated at 0.6 and 0.8 for the SSX and SFX structure, respectively. For all oxygen-bound structures occupancies were estimated through successive cycles of refinement with REFMAC systematically varying the occupancy of the ligand and assessing B-factors and electron density. An example of this occupancy determination strategy is shown in Figure S7 for the oxy-hhMb SSX structure exposed to atmospheric oxygen (Figure S6a), where the final occupancy (0.6) was chosen based on two criteria: the decrease of the B factors of the two oxygen atoms – approaching B factor values of the nearby residues and the heme – and the disappearance of difference density (Fo-Fc contoured at 3σ) close to O_2_.

**Figure 2.**
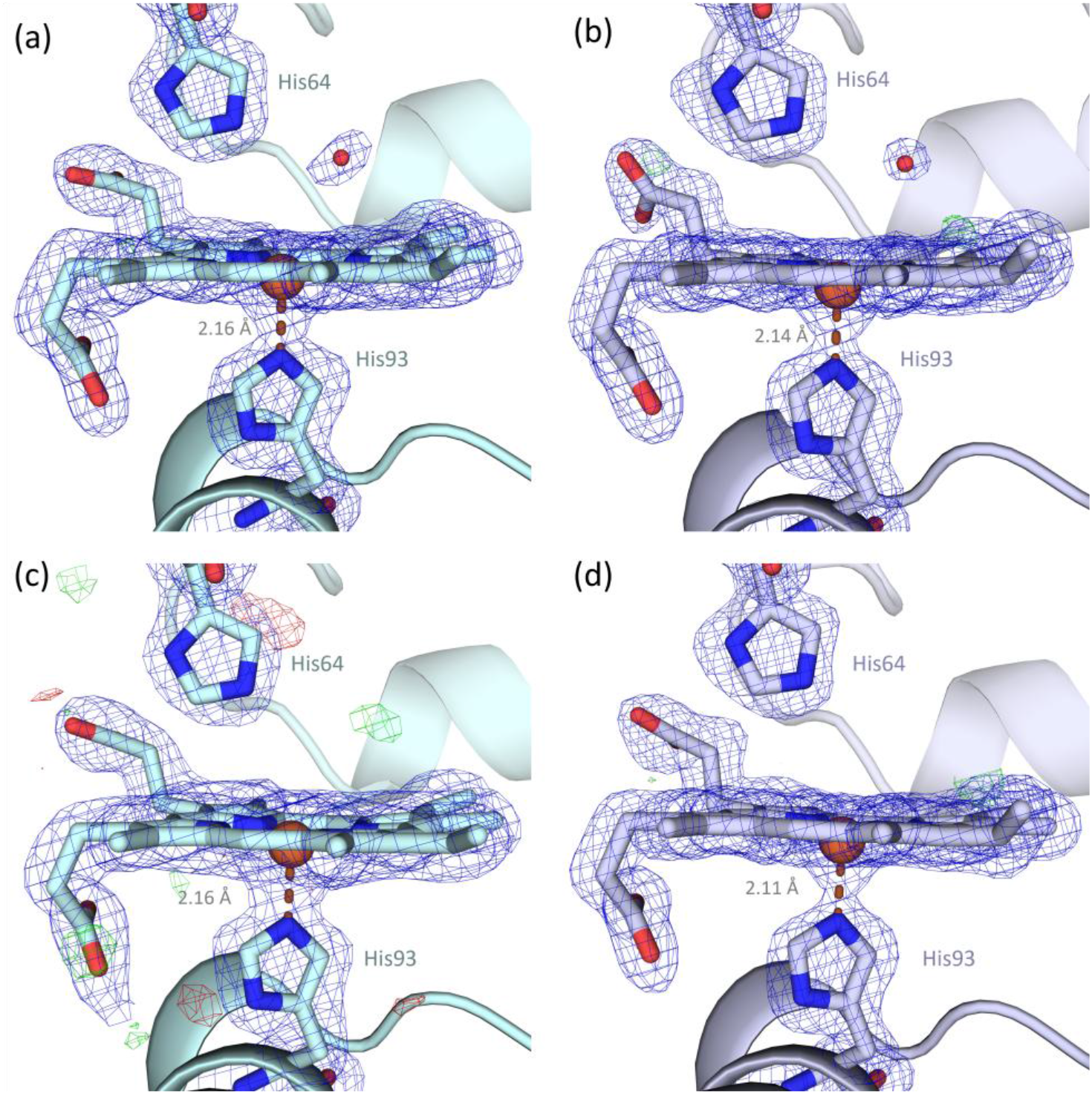
Deoxy hhMb (a) SSX (1.8 Å resolution) and (b) SFX (1.7 Å resolution) structures collected at room temperature from sodium dithionite treated microcrystals on silicon fixed targets under fully anaerobic conditions. (c) Control SSX structure (2.0 Å resolution) collected from reduced microcrystals soaked in the O_2_ cage. (d) Control SFX structure (1.7 Å resolution) from reduced microcrystals soaked in the O_2_ cage with a probe-pump collection scheme, where the pump laser was activated after the X-ray pulse as detailed in the text. 2Fo-Fc and Fo-Fc maps contoured at 1 and ±3σ, respectively.

The continued anaerobicity of samples over prolonged time periods was also confirmed through collection of *in situ* UV-Vis spectra of reduced hhMb as shown in Figure S4. hhMb microcrystals were reduced in a glovebox and loaded onto a “chipless” chip (*i.e.* between two layers of EVAL without the silicon fixed target chip) as shown in Figure S4a. *In crystallo* UV-Vis spectra of reduced hhMb microcrystals enveloped with EVAL were collected using an offline microspectrometer (Figure S4b). Tracking the Q band region between 500 – 600 nm, where a double peak indicates oxygenated myoglobin, no change was observed that could be associated with O_2_ penetration through the EVAL film (Figure S4c).

The combination of crystallographic and spectroscopic approaches allowed iterative improvement of sample loading, chip sealing and transfer resulted in the optimised workflow for anaerobic data collection, described in detail in Methods. Key aspects of this workflow are the use of single layers of EVAL on each side of the chips as a robust O_2_ barrier film, and the preparation of hhMb in the presence of excess sodium dithionite to ensure fully reduced conditions. Although an excess concentration of dithionite was used as a precaution to guarantee complete reduction across the crystal population, spectroscopic measurements indicate that substantially lower concentrations (50 mM) of dithionite are sufficient to fully reduce hhmb microcrystals (Figure S5b), and the reduced state remains stable when crystals are protected from an aerobic atmosphere with EVAL film.

In deoxy hhMb, the heme iron is in the ferrous Fe (II) state sitting slightly out of the porphyrin plane towards the proximal histidine ligand (His93). The out-of-plane iron distance is 0.29 and 0.33 Å for the deoxy-hhMb SSX and SFX structures, respectively, as shown in Table S2. The Fe-N(His93) distance was observed to be 2.16 Å and 2.14 Å for the deoxy hhMb structures in Figures 2a and 2b, respectively. Upon O_2_ binding, the effective radius of the iron atom decreases, moving the iron into the porphyrin plane pulling the proximal histidine, while reducing heme doming. This is reflected in a shortening of the Fe-His93 distance to 2.05 Å and 2.08 Å in the oxy structures (Figure S6a and S6b). All interatomic distances between the iron and His93 and His64, as well as between iron and the ligand (O1 atom) - where applicable - and out of plane distances are summarised in Table S2.

We note that there are no clear X-ray induced differences between the ultra-low dose (less than 1 kGy) SSX structure and that obtained via SFX at SACLA (Figure 2a and 2b). Low dose SSX data was facilitated by data collection at 20 keV and use of a CdTe-based detector. This strategy increases the information per unit dose that can be obtained in both conventional and serial experiments (*52*).

### Filtering data with xia2.ssx

In addition to the expected improvement in merging statistics, ΔCC_1/2_ filtering in DIALS at the scaling step was found to significantly, and concomitantly, improve the quality of electron density maps and refinement statistics. The filtered dataset produces substantially better-defined electron density throughout the structure, particularly within the heme pocket (Figure 1c). The density of the distal water molecule exhibits a clear, spherical shape whereas the corresponding density in the unfiltered dataset is less well resolved. Positive difference density peaks, that can’t be easily modelled, as the green density peak fusing into the heme density (Figure 1c) are also apparent in the unfiltered dataset. These hard to model difference map noise peaks were also observed in the vicinity of other amino acids across the model. Both scaling and refinement statistics are significantly improved when ΔCC_1/2_ filtering is applied, reducing the R_split_ value by a factor of two, improving the Wilson plot (Figure S9), and significantly decreasing R_work_ and R_free_ (Table 2). To ensure a fair comparison and minimise potential sources of bias, both datasets were refined using identical protocols using the same search model, number of refinement cycles, and identical refinement parameters. Further evidence for the improved quality of the filtered dataset is provided by solvent modelling: 41 waters could be confidently modelled in the filtered structure compared to only 33 in the unfiltered, indicating a more interpretable and better refined model.

### Release of molecular oxygen from the photocage and control experiments

A series of control experiments were conducted to verify that the caged O_2_ compound remained intact under the reducing conditions used for serial data collection, and to ensure that decaging and subsequent binding of O_2_ to hhMb did not occur without direct photoexcitation. These controls were particularly important considering previous reports (*17*) suggesting that the cage may undergo degradation in the presence of dithionite, potentially leading to unintended O_2_ release.

For the control experiments, deoxy hhMb crystals were soaked in 10 mM cage solution after removal of the microcrystalline slurry supernatant. Before data collection, only a small volume, sufficient for loading 1-2 chips, was prepared to avoid long soaking times and potential cage degradation. Two different types of control experiments were performed under fully anaerobic conditions: data were measured from cage-soaked crystals at I24 without using any laser photoexcitation to detect any potential O_2_ release from the cage resulting from the local chemical environment (dithionite). The second control experiment was performed in probe-pump format at SACLA where the 308 nm pump laser was inadvertently triggered 10 ms after collecting diffraction data (*i.e.* diffraction data were collected from chip aperture *n* immediately after aperture *(n-1)* was photo-excited) rather than as in ‘standard’ pump-probe as intended. The respective SSX and SFX control structures of hhMb are shown in Figure 2c and 2d, clearly representing a deoxy state. The absence of O_2_ binding is evident in these two structures, providing strong structural evidence that the O_2_ photocage is not chemically degraded due to the reducing conditions and O_2_ is not spontaneously released without direct illumination with the 308 nm pump laser. Moreover, interatomic distances between the heme iron and His93 of 2.15 Å and 2.16 Å for SSX and SFX controls respectively (Table S2) signify that the iron is still in its ferrous (Fe^2+^) oxidation state, and slightly pulled out of the porphyrin plane, therefore being nearly identical to the respective values for the deoxy structures in Figure 2. The iron out-of-plane distances for the control SSX and SFX structures are 0.32 and 0.33 Å, respectively (Table S2).

In deoxy hhMb, the sixth-coordination position on the distal face of the heme is vacant as expected for a ligand-free state. The distal heme pocket may contain one or more solvent molecules, typically water molecules hydrogen-bonded to the distal His64. However, these species are frequently mobile and weakly ordered leaving the heme pocket effectively empty as seen in structure Figure 2d or are only partially occupied especially in crystal forms, as seen in Figure 2c where a positive Fo-Fc difference density peak (contoured at +3σ) is clearly visible. However, the density is insufficiently defined to allow reliable modelling of a water molecule, indicating extremely low occupancy of the water molecule.

Photoactivation of the caged O_2_ compound with a 308 nm laser enabled direct observation of oxygen binding to myoglobin by time-resolved SFX at a 10 ms time delay at two different excitation energies. The O_2_-bound structures are shown in Figure 3a (28.6 µJ laser energy) and 3b (191 µJ). In both datasets, positive difference density peak (contoured at +3σ) was observed at the sixth coordination position of the heme iron, consistent with partial formation of the oxy-hhMb state following successful decaging. Isomorphous difference maps (Fo^10ms_28.6µJ^ – Fo^dark^, Figure 3c) between the 28.6 µJ dataset and the deoxy SFX (dark) state and (Fo^10ms_191µJ^ – Fo^dark^, Figure 3d) between the 191 µJ dataset and the deoxy SFX state confirm successful photo-release of O_2_ from the cage compound visualised by the positive (coloured green in Figure 3) difference density peak above the heme iron, contoured at +3σ. A negative (coloured red) difference density peak (−3σ) below the iron indicates the displacement of the iron towards the plane of the porphyrin ring upon ligand binding. Occupancy refinement, carried out as described above, yielded oxygen occupancies of approximately 0.2 when a laser energy of 28.6 µJ was used (illustrated in Figure S8), increasing to 0.35 with an incident laser energy of 191 µJ.

**Figure 3.**
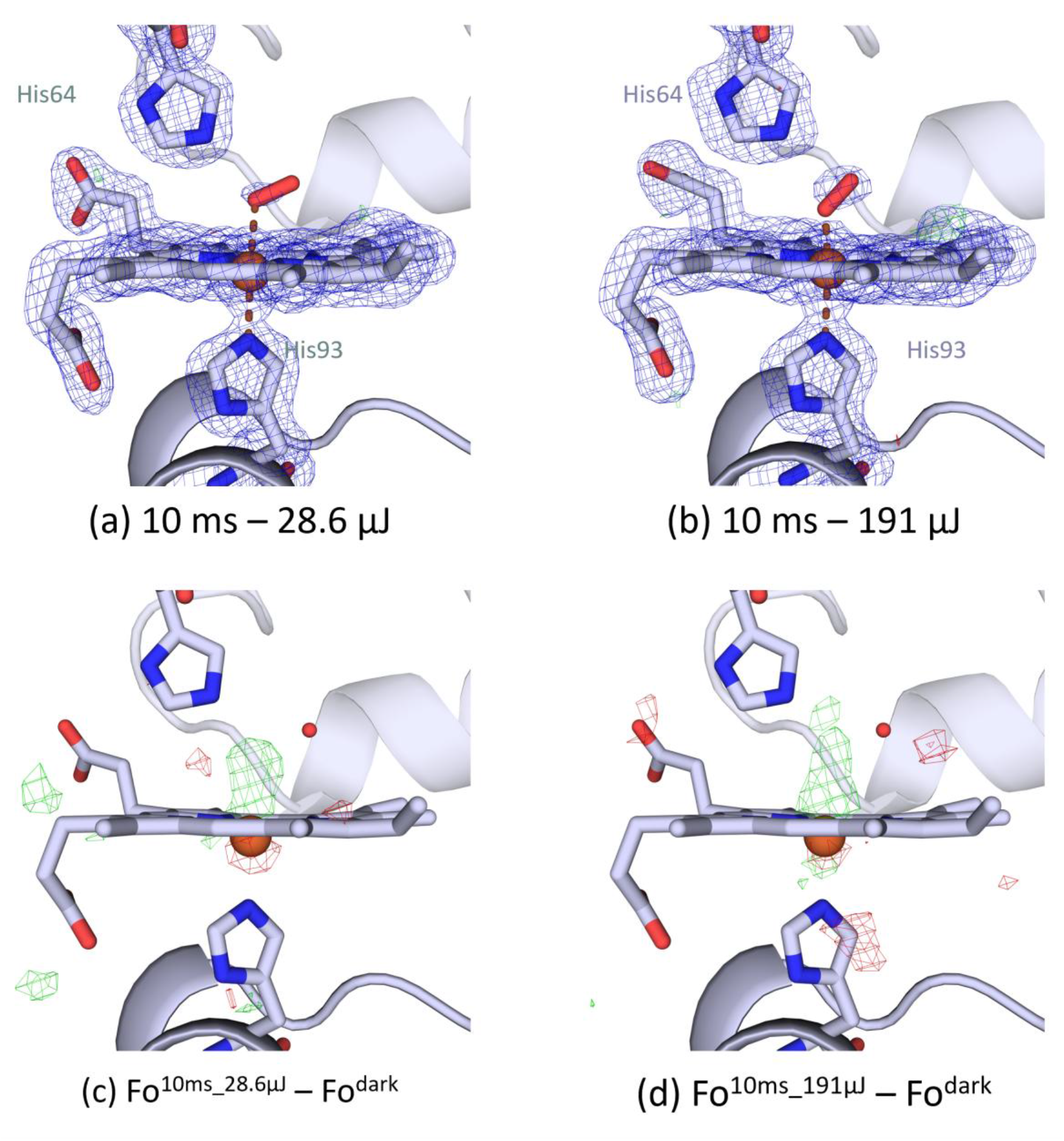
Oxygen-bound SFX hhMb structures (1.7 Å resolution) obtained using the O_2_ photocage 10 ms after a pump laser pulse using laser energies of 28.6 µJ (a) and 191 µJ (b). Oxygen occupancy was estimated at 0.2 and 0.35, respectively. 2Fo-Fc and Fo-Fc maps contoured at 1 and ±3σ, respectively. Isomorphous difference maps (c) Fo^10ms_28.6µJ^ – Fo^dark^ and (d) Fo^10ms_191µJ^ – Fo^dark^, contoured at ± 3σ, showing successful photo-release of O_2_ from the cage compound (green difference density peak above heme iron). The red difference density below the iron indicates displacement of the iron towards the plane of the porphyrin ring upon O_2_ binding.

These occupancy values are consistent with the observed electron density features and indicate increased photo-release of the caged O_2_ at the higher laser power. However, the photocage SFX structure at 191 µJ laser power should be treated as less reliable due to the low numbers of crystals merged in the final dataset (see Table 1). We note that despite the relatively modest number of crystals forming the dataset, polder maps calculated with side chains, heme or bound O_2_ omitted showed clear well-defined density indicating sufficient data had been collected (*53*). Polder maps corresponding to Figure 3a and 3b are shown in supplementary Figure S10. The refined occupancies further support the interpretation that UV laser excitation successfully releases caged O_2_, leading to partial population of the oxy-hhMb state under anaerobic room-temperature conditions. The interatomic distances between the heme iron and His93, as well as those between Fe – His64 and Fe – O1 (Table S2), are at the upper end of the range typically encountered in oxy-hhMb structures suggesting partial oxygen occupancy and/or mixed oxy-/deoxy-states in the microcrystals.

It is noteworthy that the oxy-hhMb oxygen occupancies obtained by photoactivation of the cage O_2_ compound are somewhat lower than those observed in the corresponding oxidised structures (Figures S6 and S7) obtained following accidental exposure to atmospheric oxygen. This reduced occupancy is expected, as only a fraction of the caged molecules undergo photolysis upon laser excitation limiting the amount of O_2_ released during a single pump-probe experiment.

### Checkerboard data collection scheme for SSX and SFX using the oxygen photocage

To assess the extent of O_2_ release and diffusion upon decaging during time-resolved experiments, a checkerboard data collection scheme was used. This approach was implemented both at I24 and SACLA leading to some contrasting results. SSX checkboard structures are shown in Figure 4a (light), 4b (dark) and 4c (isomorphous difference map between the light and the dark datasets of the checkerboard), revealed successful decaging of the O_2_ photocage at 5 ms time delay and apparent diffusion of the ligand to the wells comprising the dark dataset. Oxygen occupancy in the illuminated wells was approximately 0.85, whereas the corresponding occupancy in the dark wells was substantially lower, reaching 0.45. The presence of significant oxygen density in both subsets suggests gas diffusion between neighbouring wells, resulting in cross-contamination across the checkerboard pattern. These observations indicate that O_2_ released in one region of the chip can migrate sufficiently over the experimental timescale to populate adjacent, nominally dark, positions blurring the claimed timepoints.

**Figure 4.**
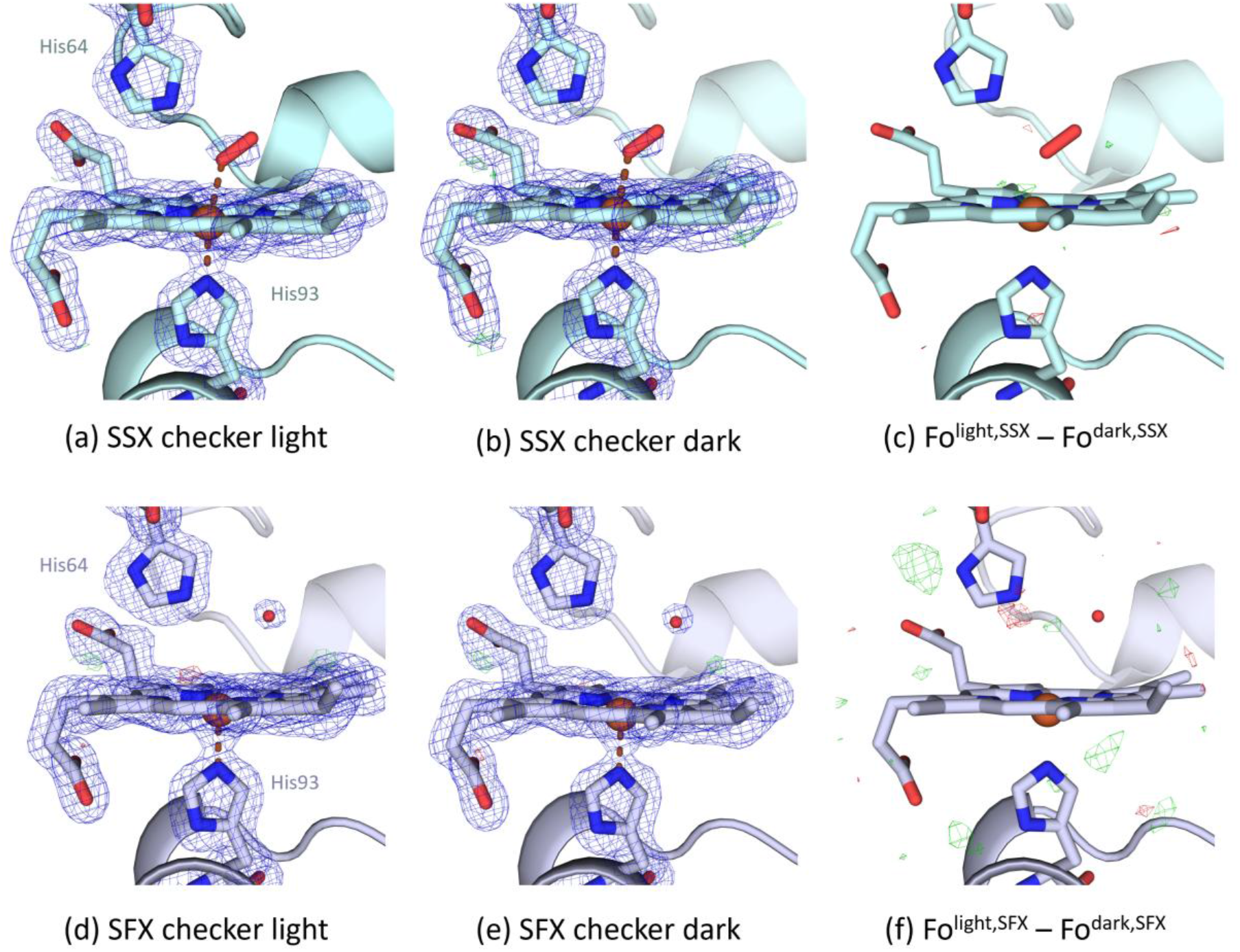
Checkerboard data collection scheme. SSX hhMb structures (1.8 Å resolution) showing (a) successful release and binding of caged O_2_ at 5 ms time delay using 25×0.47 µJ laser pulses (light dataset) and (b) the dark dataset showing cross-contamination between adjacent wells on the fixed targets. Estimated occupancies are 0.85 and 0.45, respectively. (c) Isomorphous difference map Fo^light,SSX^ – Fo^dark,SSX^ showing no major differences between the two datasets. SFX hhMb structures (1.7 Å resolution) showing (d) unsuccessful binding of caged O_2_ at 100 µs and 35 µJ laser energy and (e) the respective dark dataset of the checkboard scheme. (f) Fo^light,SFX^ – Fo^dark,SFX^ showing no difference density peak above the heme iron between the two checkerboard datasets suggesting unsuccessful O_2_ binding. 2Fo-Fc maps contoured at 1σ and Fo-Fc and Fo-Fo maps contoured at ±3σ.

In contrast, no evidence of O_2_ binding was observed in the 100 μs time-delay SFX experiment. The light and dark datasets from the checkerboard scheme are shown in Figure 4d and 4e, respectively, while the isomorphous difference map (Fo^light,SFX^ – Fo^dark,SFX^) in Figure 4f shows no apparent difference peak that could justify potential ligand binding. However, lack of bound O_2_ at the heme site in the light dataset doesn’t necessarily imply unsuccessful decaging. We previously demonstrated that a pulse energy of 28.6 µJ of the pump laser at SACLA led to decaging and subsequent O_2_ binding for the 10 ms structures shown in Figure 3a. For the checkerboard light structure (Figure 4d) a slightly higher pulse energy was used (35 µJ) at a faster time delay (100 µs). These findings suggest that O_2_ binding to myoglobin is not detectable on this short timescale under the present experimental conditions. Although photolysis of caged O_2_ has been reported to occur on sub-microsecond timescales (< 1 µs for HPBC; (*2*)) though several processes following decaging may limit observation of O_2_ binding at 100 μs, including transit of the released O_2_ through the liquid phase and across the liquid-gas interface, diffusion through crystal solvent channels, and migration to the heme pocket. The relative contribution of these process remains unclear and requires further investigation. The checkerboard data collection scheme provides improved temporal precision and may facilitate future studies aimed at disentangling uncaging kinetics from oxygen diffusion and ligand-binding events.

The stability of the photocage in crystals over timescales of hours is illustrated by data collection from hhMb microcrystals soaked in the caged O_2_ solution for approximately 8 h following an unscheduled beam interruption. Analysis of these datasets (data not shown) revealed clear O_2_ density above the heme iron in both the nominally dark and light subsets of the checkerboard pattern using the same data collection parameters (35 μJ pulse, 100 μs prior to X-ray pulse) used above. The presence of bound O_2_ in the dark dataset, despite the absence of UV excitation, indicates that O_2_ had been released from the cage prior to data collection. This observation is consistent with gradual degradation of the O_2_ photocage during prolonged incubation in the presence of sodium dithionite. Our findings suggest that although prolonged exposure to sodium dithionite can ultimately compromise photocage stability, this was not observed for datasets where deoxy-hhMb crystals were soaked in cage for < 1 h, suggesting that the half time of ∼ 1 s for the O_2_ cage degradation in sodium dithionite determined in solution by (*17*) may not apply in crystals.

## Conclusions/discussion

The results presented above establish a robust and experimentally straightforward approach for performing anaerobic room-temperature serial crystallography using fixed targets at both synchrotron and XFEL facilities. By combining low oxygen-permeability EVAL film with a robust sample preparation workflow, anaerobic conditions can be reliably maintained over the timescales required to collect diffraction data. The use of a single EVAL film layer on each side of the sample holder simplifies sample preparation making it essentially identical to ‘standard’ fixed target data collection (*25*) while retaining excellent protection from atmospheric oxygen. Mb readily oxidises *in crystallo* when in contact with atmospheric oxygen, and the determination of anaerobic room-temperature SSX/SFX structures of deoxy hhMb illustrates the effectiveness of the approach. The methodology can be readily applied to crystals of any oxygen-sensitive system.

Furthermore, the approach is well suited to the use of photocages. While caged oxygen compounds have previously been used extensively in spectroscopic studies of oxygen-dependent enzymes, direct structural demonstration of controlled photo-released O_2_ binding has remained elusive. Here, oxygen released by UV excitation of the HPBC cage was directly visualised through binding to hhMb on the millisecond timescale. No evidence of oxygen occupancy was observed in samples mixed with photocage but not exposed to laser light, demonstrating that the cage remains stable during sample preparation and data collection and that O_2_ release is triggered on demand by laser illumination. It is noteworthy that full O_2_ occupancies were not observed with the photocage, supporting the conclusion that oxygen release is directly controlled by photoexcitation rather than inadvertent exposure to atmosphere. Reduced occupancies highlight possible future developments as high populations of reaction intermediates are typically required to generate interpretable electron density and future experiments could benefit from cages with higher quantum yields and/or greater solubility. The absence of detectable O_2_ binding in the 100 μs SACLA data indicate that under the conditions used, O_2_ release from the cage and subsequent binding to hhMb is not observed on this timescale.

## Acknowledgements

We gratefully acknowledge the BBSRC (BB/W001950/1 and BB/X01844X/1) for supporting S.J., A.L. de la I., M.L., J.A.R.W., R.L.O., M.A.H. XFEL experiments were performed at BL2 EH3 of SACLA with the approval of the Japan Synchrotron Radiation Research Institute (JASRI) (proposal numbers 2022B8041, 2023B8017, 2024B8003 and 2025B8021). We acknowledge use of Diamond beamline I24 under proposals MX28583 and MX36015. The PORTO laser at Diamond was funded by the EPSRC (EP/P001548/1).

## Supplementary information

### S1. Spectroscopic validation of the oxygen photocage synthesis

**Figure S1.**
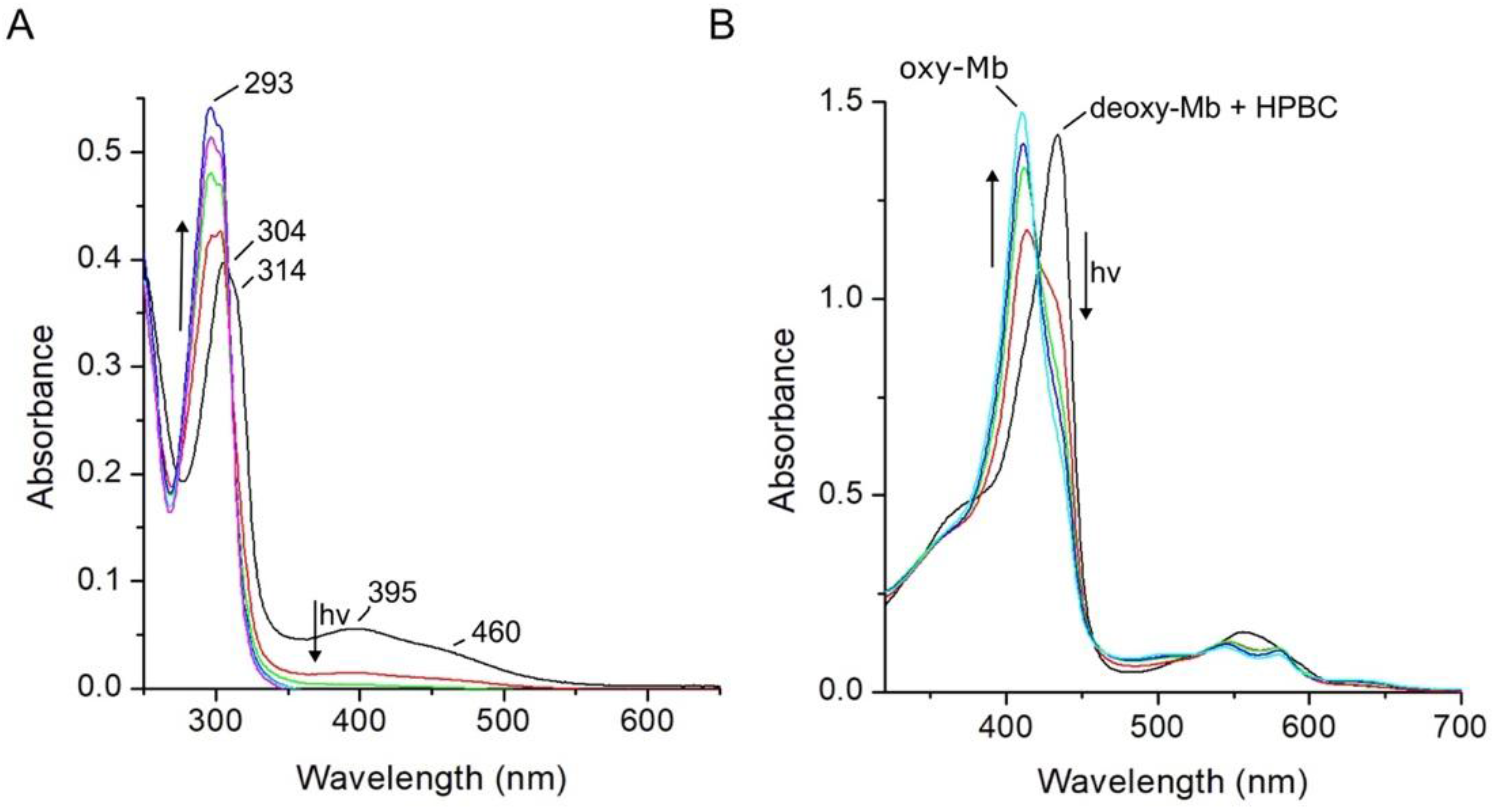
Solution UV-Vis spectroscopy of (A) the synthesised HPBC perchlorate salt and (B) hhMb solution mixed with HPBC under anaerobic conditions at pH 7. The direction of spectral change following white light flash and decaging (*i.e.* O_2_ release) is indicated by the arrows. The wavelength maxima and spectral changes before (460, 395, 314, 304 nm) and after (293 nm) continuous white light flashing of the HPBC complex in (A) are consistent with published data (*15*). In (B) continuous white light flashing of the starting deoxy-hhMb sample mixed with HPBC initiates a spectral change consistent with O_2_ release from HPBC and formation of oxy-hhMb.

### S2. Validation of anaerobic data collection methods

**Figure S2.**
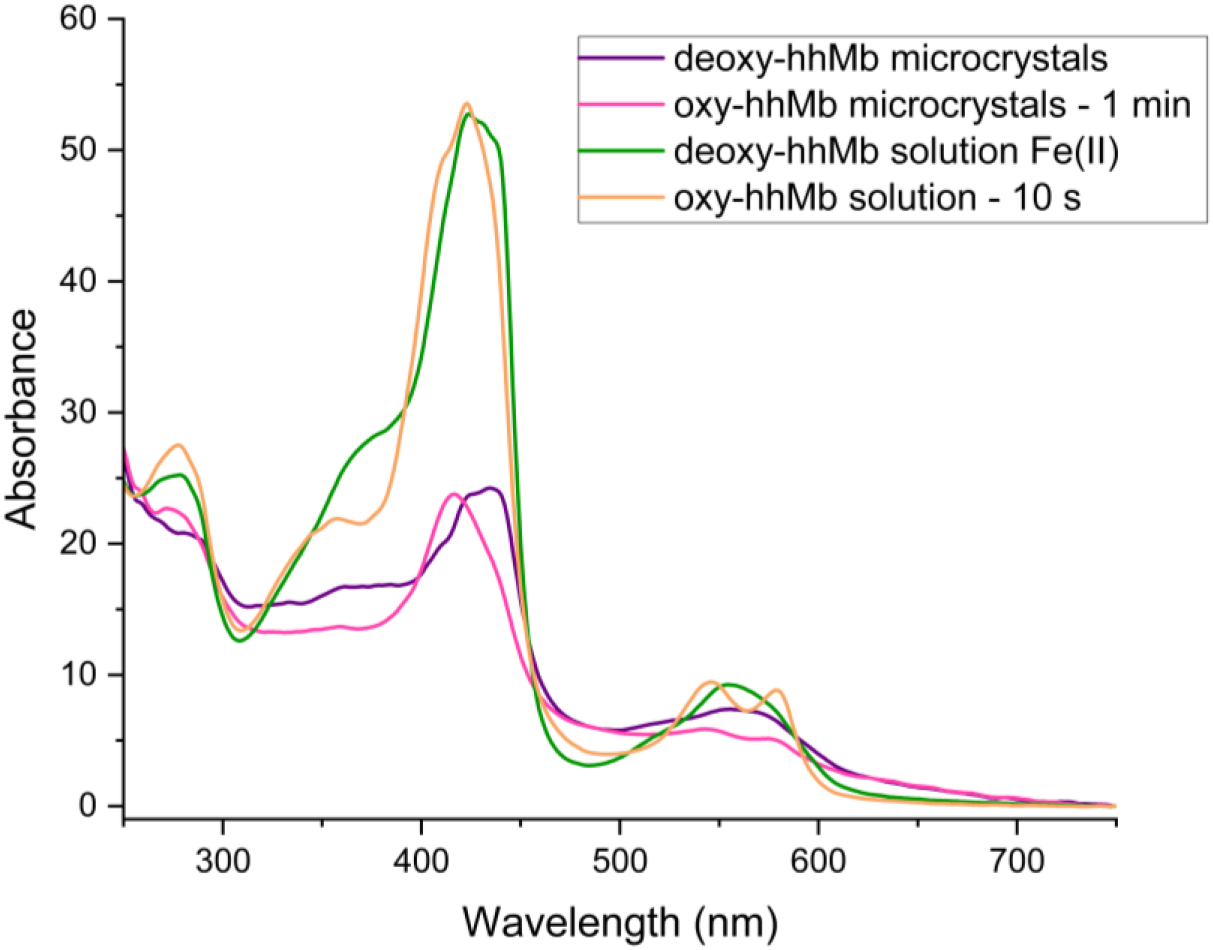
Solution and microcrystal spectra of deoxy-(purple/microcrystals, green/solution) and oxy-(pink/microcrystals, orange/solution) hhMb by exposure to atmospheric oxygen. Spectra of reduced samples were collected within an anaerobic chamber. Data were collected using a DS11 spectrometer (DeNovix).

**Figure S3.**
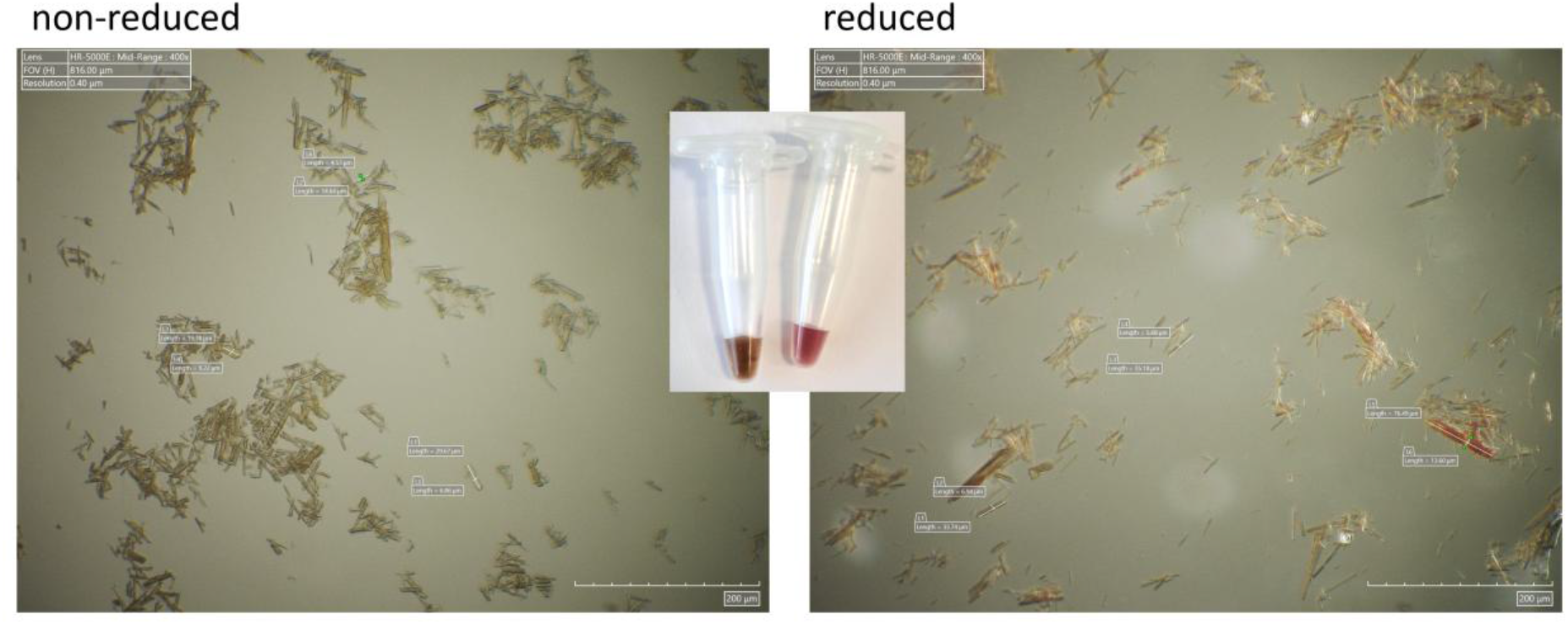
Images of oxidised (left) and reduced (right) hhMb crystals taken using a high-resolution Hirox microscope. Myoglobin tends to form relatively long needles or thin plates forming clusters, necessitating thorough mixing of the crystalline slurry prior to loading the samples onto the fixed targets. Here, microcrystals were reduced in the presence of excess sodium dithionite and the characteristic colour change from brown (oxidised) to bright red (reduced) is evident from the microscope images and the Eppendorf tubes containing the microcrystals (insert). Scale bar represents 200 µm.

UV-Vis absorption spectra (Figures S4 and S5) were recorded using a microspectrophotometer developed at beamline I24 for on- and off-line *in crystallo* optical spectroscopy. The setup features two off-axis reflective objectives, a Shamrock 303i (Andor Technology) spectrometer, a Newton EM CCD detector and a fibre-coupled Xenon light source (Thorlabs) with a continuous spectrum over the range of 250–800 nm. 1 µL of hhMb crystals, either in the reduced or oxidised state, were deposited between two single layers of 12.5 µm EVAL (Kuraray) film in a “chipless” chip format and mounted at the focal point of the two objectives. The spectra were an accumulation of 5-10 exposures each of 10 ms duration. The focal spot of the white light was approximately 50 μm. The spectra shown in Figures S4b and S4c were processed and plotted with an in-house Python script using a Savitzky–Golay filter for smoothing while the baseline was corrected using the ModPoly method of the BaselineRemoval library. The spectra shown in Figure S5 were processed and plotted with OriginPro using a Savitzky-Golay filter for denoising.

**Figure S4.**
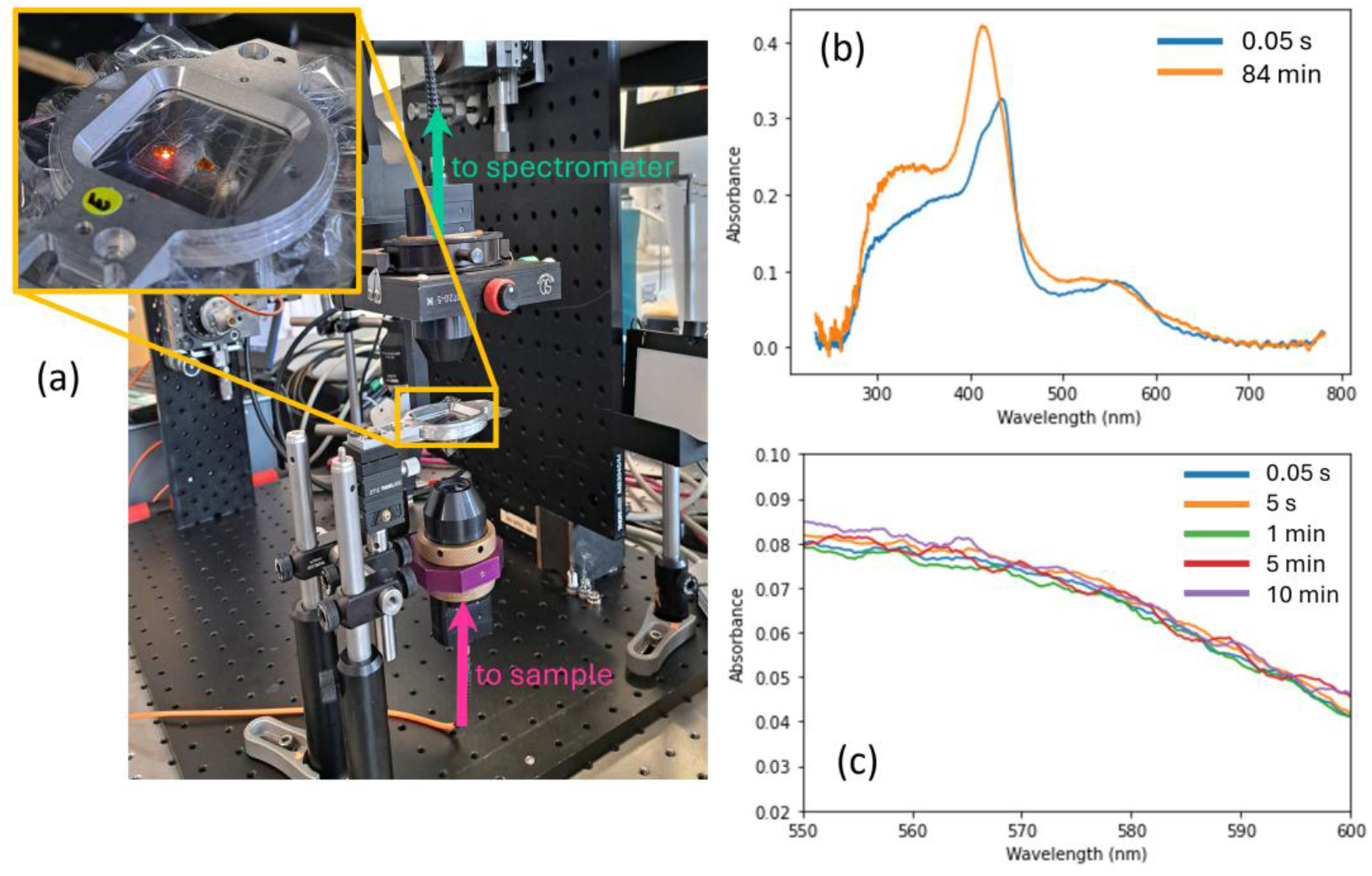
Offline setup for recording spectra from crystals and solutions in fixed targets and thin films (a). Spectra obtained from hhMb crystals held between two EVAL films over 84 mins (b) and over Q-band region over approximate timescale for fixed target data collection (c).

The UV-Vis spectra below show that hhMb microcrystals can be successfully reduced in 50 mM sodium dithionite solution and there is no need to continue working in excess quantities of the reducing agent. During early experiments when the protocols for robust anaerobic data collections were developed, a choice was made to work under excess quantities of dithionite ensuring that the sample would be thoroughly reduced in the microcrystal slurry and would stay reduced for the whole duration of the experiment. The respective spectra recorded under excess dithionite conditions are shown in Figure 1b.

**Figure S5.**
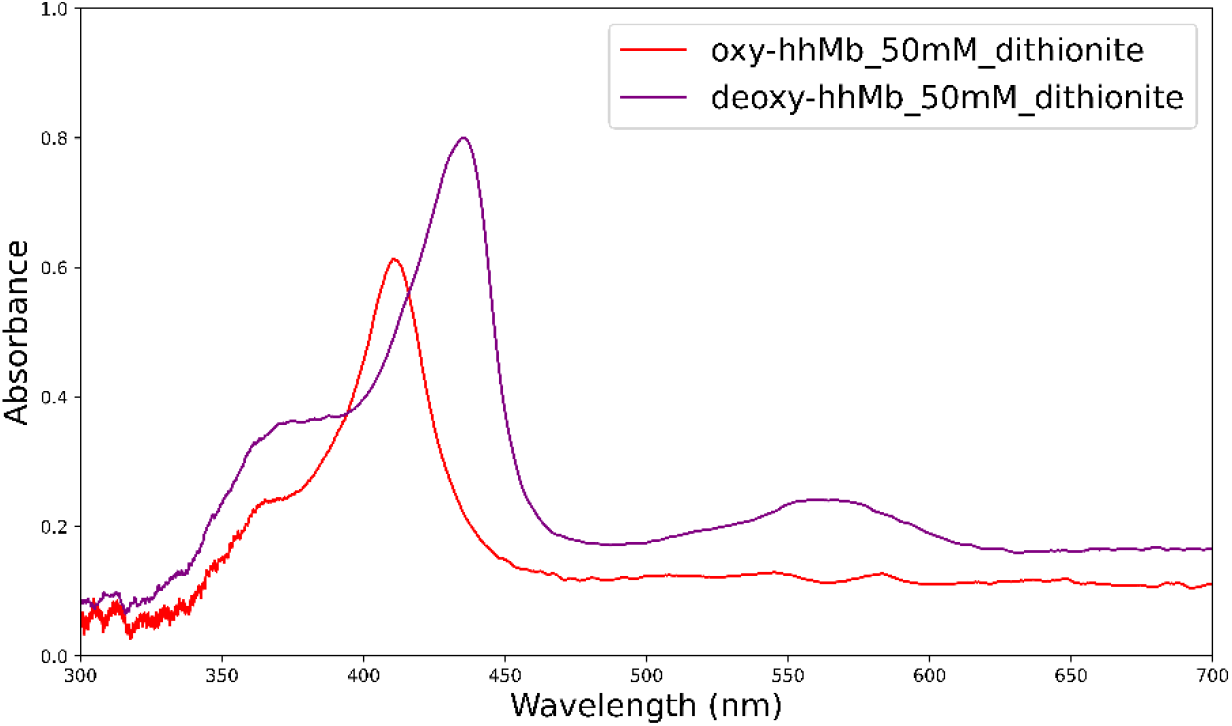
UV-Vis spectra obtained from hhMb microcrystal slurries on “chipless” chips of EVAL film. Reduced (Fe^2+^) and oxidised (Fe^3+^) hhMb microcrystals in 50 mM dithionite (purple and red lines respectively). Reduction of hhMb crystals took place in a glovebox. 1 µL of the microcrystalline slurry was transferred onto each of the wells of an adhesive tape placed above one layer of EVAL. A second layer of EVAL would encapsulate the droplets in these “chipless” chips and the chip holder was ported out of the glovebox for subsequent spectroscopic measurements. For the oxidised measurements, the “chipless” chips were instantly opened in atmospheric oxygen and resealed again.

### S3. Oxidised SSX and SFX structures

Results of early experiments trying to validate the anaerobic environment and to capture room-temperature reduced (dark) structures of hhMb with serial X-ray crystallography are shown in Figure S6. These are two oxidised structures including one collected at I24 (Figure S6a) where the crystals were not properly reduced and one collected at SACLA XFEL (Figure S6b) where the glovebox was accidentally opened during the experiment. The heme site view shows evident oxygen binding to the iron with the occupancies estimated at 0.6 (Figure S7) and 0.8 (not shown) for the SSX and the SFX dataset, respectively. The data processing and refinement statistics are given in Table S1. The interatomic distances between the iron and proximal (His 93) and distal (His 64) histidine and the oxygen ligand are given in Table S2, along with the respective distances for the deoxy SSX and SFX structures shown in Figure 1.

**Figure S6.**
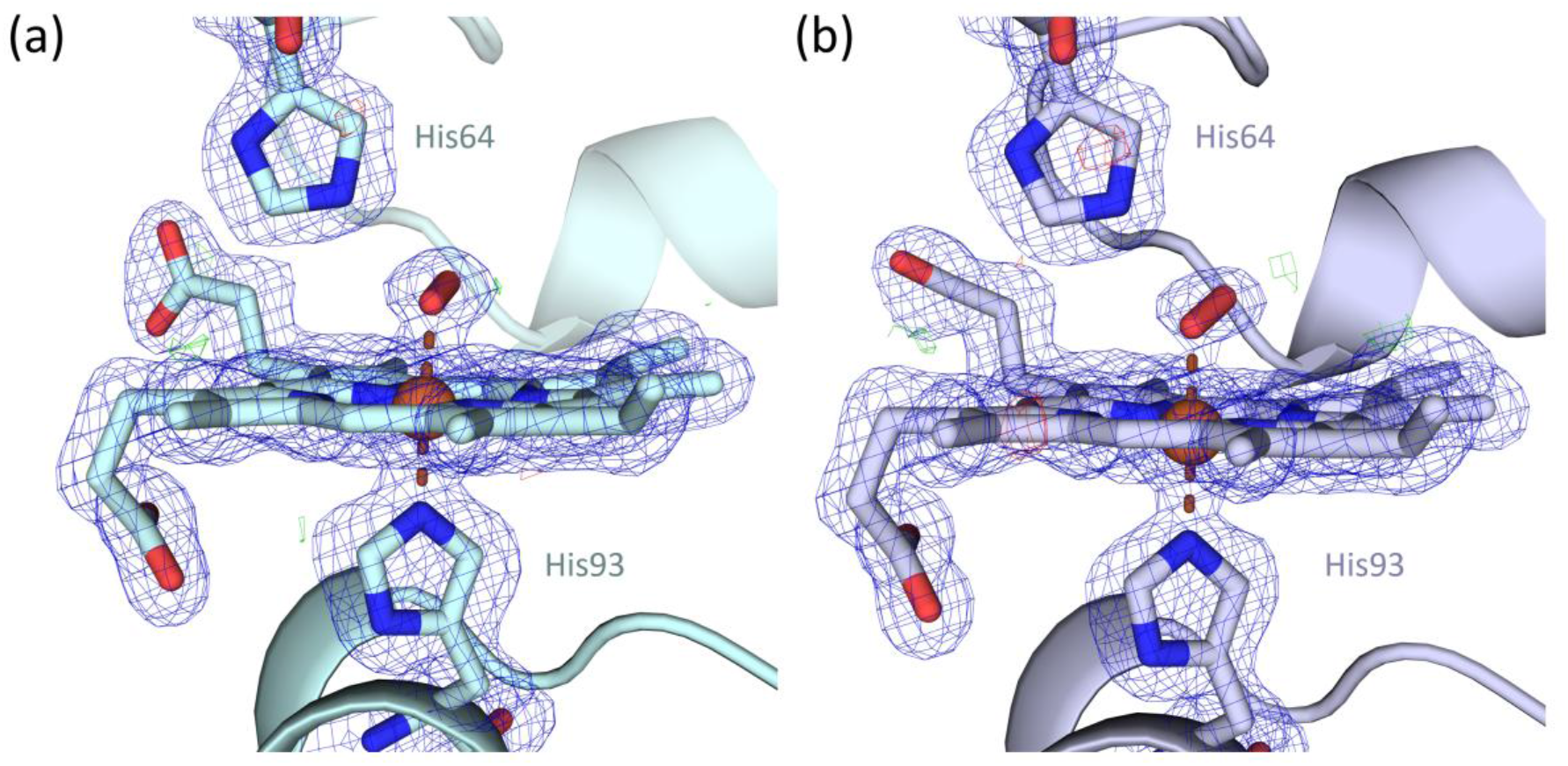
Oxy-hhMb (a) SSX and (b) SFX structures (both at 1.7 Å resolution) of reduced crystals accidentally exposed to atmospheric oxygen. 2Fo-Fc and Fo-Fc maps contoured at 1 and ±3σ, respectively.

**Table S1.** Data processing and refinement statistics for oxy-hhMb SSX and SFX structure exposed to atmospheric oxygen. Values in parentheses refer to the outermost resolution shell. For all data the space-group was P12_1_1. The SSX structure was indexed, integrated and scaled using xia2.ssx (*33*) and the SFX structure using CrystFEL 0.11.1 (*34*).

| Dataset Name | Oxy-hhMb SSX<br>(atmospheric exposure) | Oxy-hhMb SFX<br>(atmospheric exposure) |
| --- | --- | --- |
| X-ray source | I24, DLS | SACLA |
| X-ray energy / keV | 12.4 | 11.0 |
| Processing | xia2.ssx | CrystFEL |
| Resolution (Å) | 61.45-1.73 (1.76 - 1.73) | 61.34 - 1.67 (1.69 - 1.67) |
| Unit cell (Å) | 64.33, 28.79, 35.90<br>90, 107.22, 90 | 64.53 28.84 35.95<br>90 107.20 90 |
| Merged crystals | 6609 | 6780 |
| Total observations | 453947 (14194) | 343588 (11247) |
| Unique reflections | 13500 (677) | 15016 (774) |
| Completeness | 100 (100) | 100 (100) |
| Multiplicity | 33.6 (21.0) | 56.3 (14.5) |
| CC<sub>1/2</sub> | 0.981 (0.344) | 0.921 (0.711) |
| R<sub>split</sub> | 0.183 (1.188) | 0.171 (0.371) |
| I/σ(I) | 12.8 (1.4) | 7.4 (3.3) |
| R<sub>work</sub> | 0.178 (0.283) | 0.168 (0.241) |
| R<sub>free</sub> | 0.235 (0.328) | 0.196 (0.324) |
| RMSD bond length (Å) | 0.0128 | 0.0137 |
| RMSD bond angles (°) | 2.220 | 2.179 |
| Ramachandran favoured (%) | 94.74 | 94.74 |
| O<sub>2</sub> occupancy | 0.6 | 0.8 |
| PDB ID | pdb_000032ZA | pdb_000032YZ |

**Table S2.** Interatomic distances between iron, His 93, His 64 and O_2_ (O1 atom) for all the deoxy, oxidised by atmospheric oxygen and oxycage-bound structures reported in this work. Iron o-o-p refers to iron out-of-plane distance, with the plane defined by the four co-ordinating heme nitrogen atoms.

|  | **Fe – His93**<br>**NE2**<br>**(Å)** | **Fe – His64**<br>**NE2**<br>**(Å)** | **Fe – O1**<br>**(Å)** | **His64 NE2-**<br>**O1**<br>**(Å)** | **O-O**<br>**(Å)** | **Iron o-o-p**<br>**(Å)** |
| --- | --- | --- | --- | --- | --- | --- |
| **Deoxy SSX**<br>(Figure 2a) | 2.16 | 4.61 | n/a | n/a | n/a | 0.29 |
| **Deoxy SFX**<br>(Figure 2b) | 2.14 | 4.64 | n/a | n/a | n/a | 0.33 |
| **Oxidised SSX**<br>(Figure S6a) | 2.05 | 4.52 | 2.06 | 2.95 | 1.20 | 0.07 |
| **Oxidised SFX**<br>(Figure S6b) | 2.08 | 4.65 | 2.00 | 2.96 | 1.18 | 0.16 |
| **Control SSX**<br>(Figure 2c) | 2.15 | 4.48 | n/a | n/a | n/a | 0.32 |
| **Control SFX**<br>(Figure 2d) | 2.16 | 4.63 | n/a | n/a | n/a | 0.33 |
| **10 ms – 28.6 µJ SFX**<br>(Figure 3a) | 2.11 | 4.53 | 2.25 | 2.59 | 1.24 | 0.21 |
| **10 ms – 191 µJ SFX**<br>(Figure 3b) | 2.15 | 4.61 | 1.85 | 2.91 | 1.21 | 0.18 |
| **Checker light SSX**<br>(Figure 4a) | 2.07 | 4.61 | 2.26 | 2.89 | 1.24 | 0.18 |
| **Checker dark SSX**<br>(Figure 4b) | 2.09 | 4.58 | 2.30 | 2.85 | 1.25 | 0.18 |
| **Checker light SFX**<br>(Figure 4d) | 2.15 | 4.60 | n/a | n/a | n/a | 0.31 |
| **Checker dark SFX**<br>(Figure 4e) | 2.11 | 4.61 | n/a | n/a | n/a | 0.33 |

### S4. Occupancy estimation and refinement

Occupancy of the O_2_ bound in the oxy-hhMb SSX structure (Figure S6a) was estimated through successive refinement cycles with REFMAC by varying the occupancy value of the ligand (Figure S7). The final occupancy (0.6) was chosen based on two criteria: the decrease of the B factors of the two oxygen atoms – approaching B factor values of the nearby residues and the heme – and the disappearance of difference (Fo-Fc contoured at ±3σ) densities close to the O_2_ ligand.

**Figure S7.**
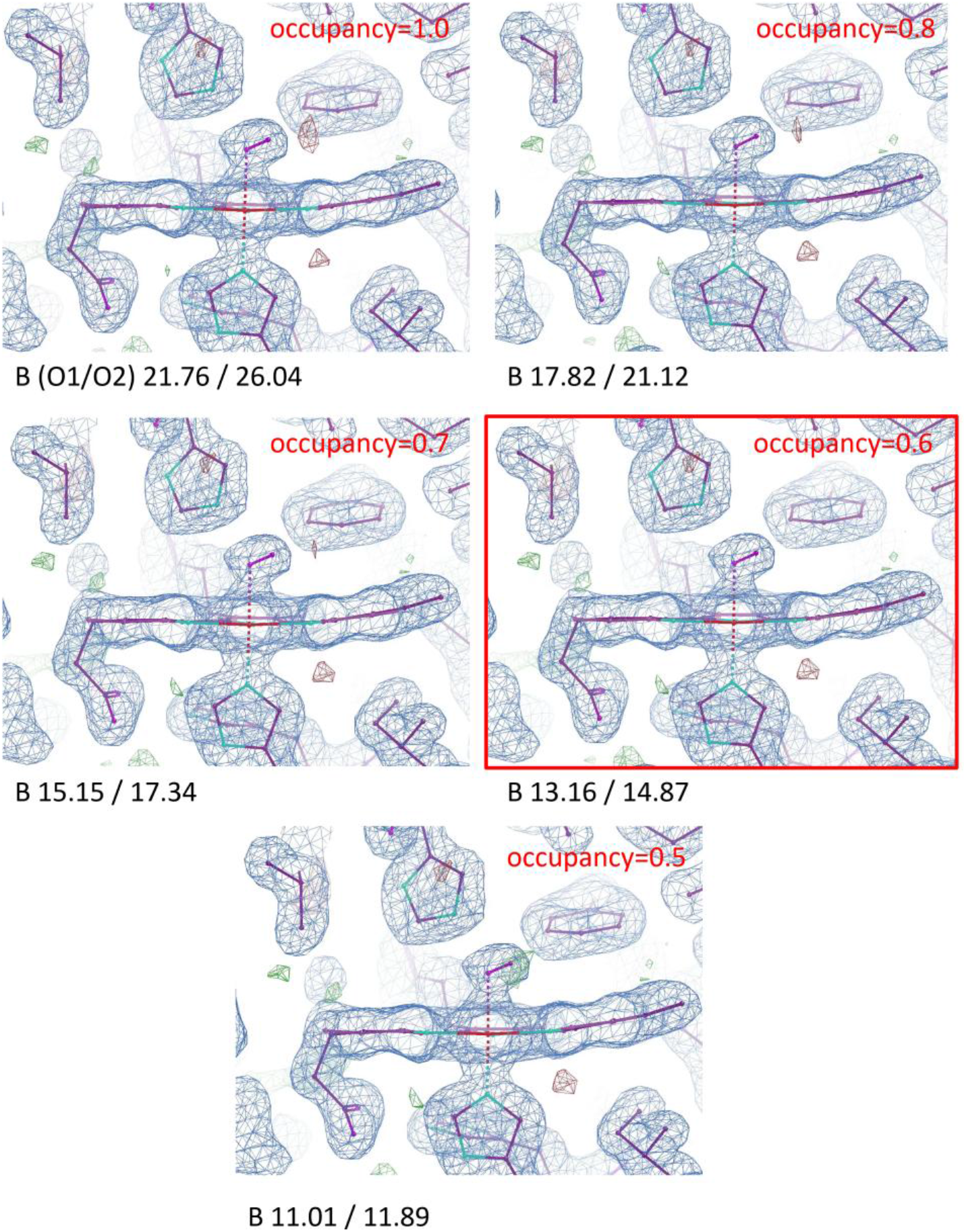
Occupancy estimation through occupancy refinements of the O_2_ bound to heme iron for the oxidised SSX structure shown in Figure S6a and B factors for O1/O2 atoms. The occupancy chosen is indicated by a red border (0.6). 2Fo-Fc and Fo-Fc maps contoured at 1 and ±3σ, respectively.

**Figure S8.**
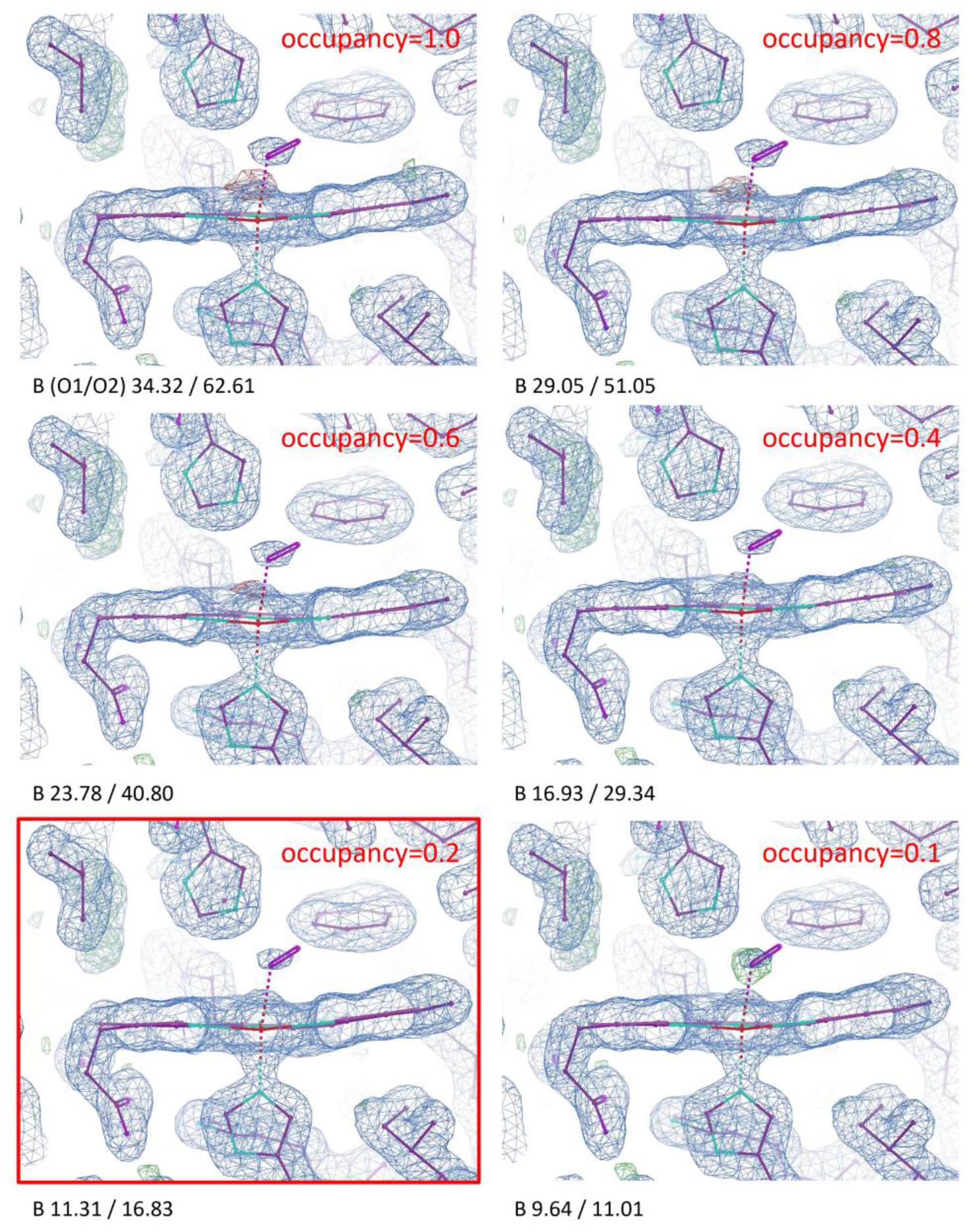
Occupancy estimation of the O_2_ for the oxygen-bound SFX structure shown in Figure 3a using an O_2_ photocage and B factors for O1/O2 atoms. Estimated occupancy at 0.2. 2Fo-Fc and Fo-Fc maps contoured at 1 and ±3σ, respectively.

### S5. Wilson plots for ΔCC1/2 filtering

**Figure S9.**
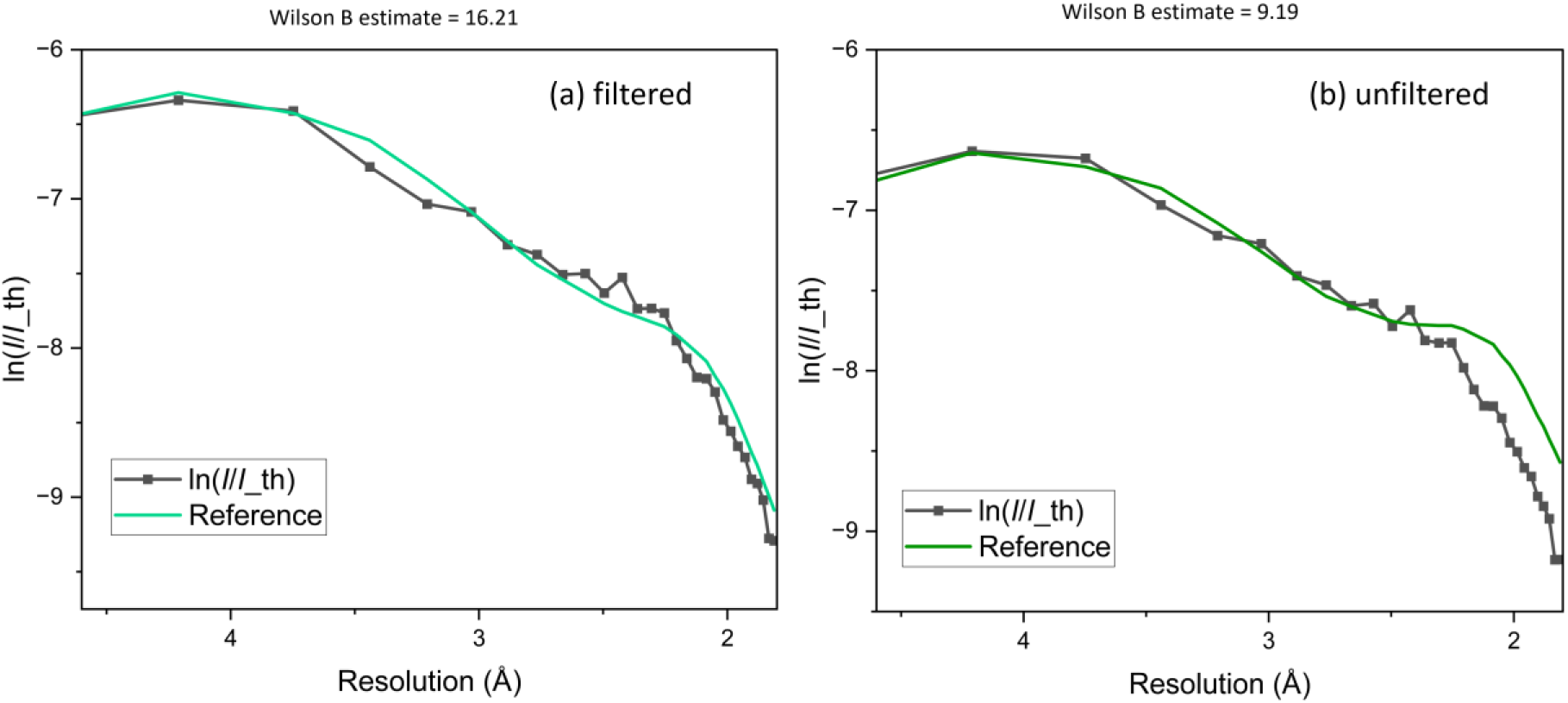
Wilson plots for the ΔCC_1/2_ filtered (a) and unfiltered (b) datasets shown in Figure 1c.

### S6. Omit maps

**Figure S10:**
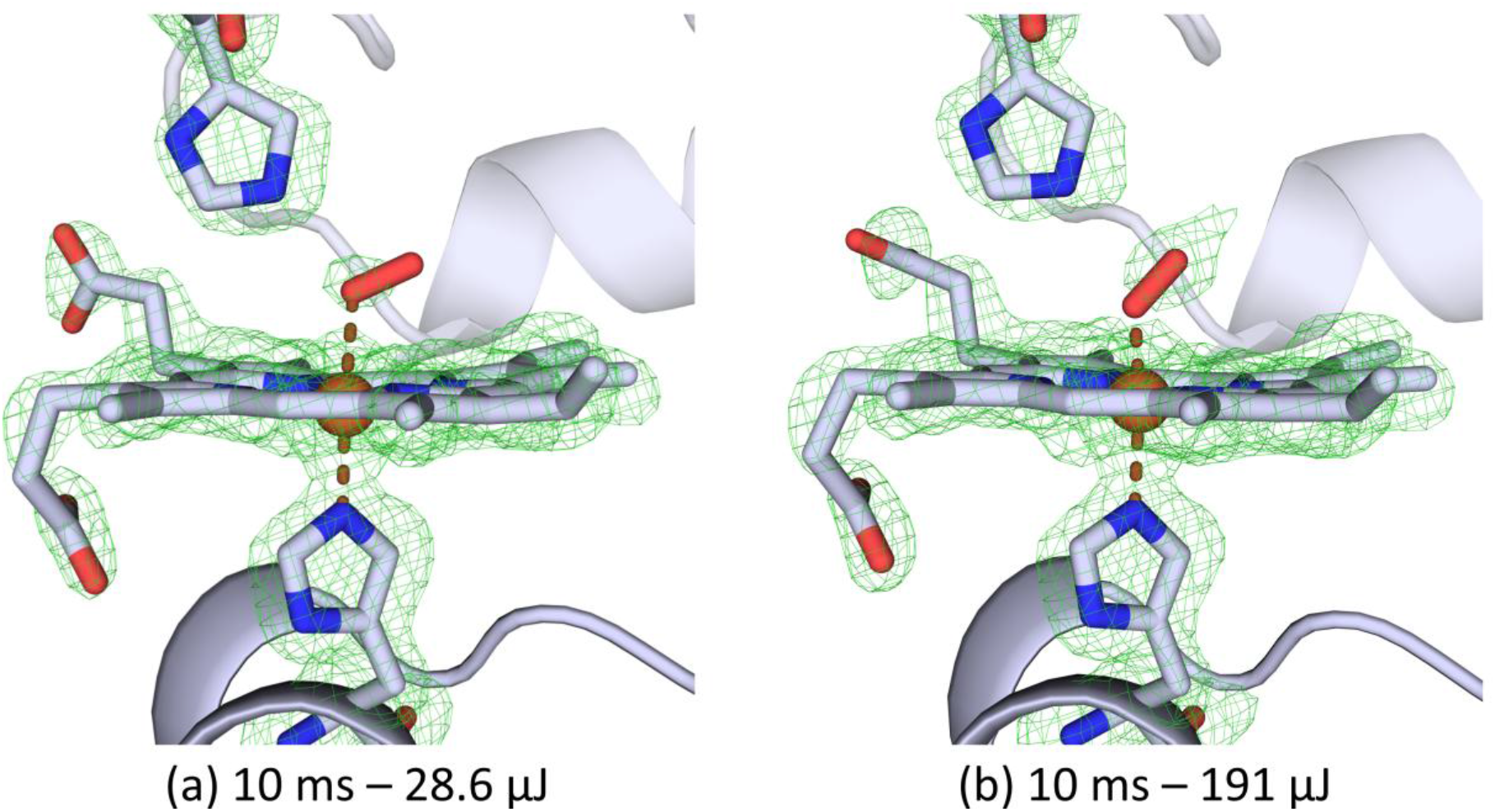
Polder omit maps corresponding to Figures 3a and 3b. Density contoured at 3σ.

## References

1. N. Caramello, A. Royant, From femtoseconds to minutes: time-resolved macromolecular crystallography at XFELs and synchrotrons. Acta Cryst D 80, 60–79 (2024).

2. D. C. F. Monteiro, E. Amoah, C. Rogers, A. R. Pearson, Using photocaging for fast time-resolved structural biology studies. Acta Cryst D 77, 1218–1232 (2021).

3. S. Kesgin-Schaefer, J. Heidemann, A. Puchert, K. Koelbel, B. A. Yorke, N. Huse, A. R. Pearson, C. Uetrecht, H. Tidow, Crystal structure of a domain-swapped photoactivatable sfGFP variant provides evidence for GFP folding pathway. The FEBS Journal 286, 2329– 2340 (2019).

4. J. I. Zaitseva-Kinneberg, A. Puchert, Y. Pfeifer, H. Yan, B. A. Yorke, H. M. Müller-Werkmeister, C. Uetrecht, J. Rehbein, N. Huse, A. R. Pearson, M. Sans, Synthesis and characterisation of α-carboxynitrobenzyl photocaged l-aspartates for applications in time-resolved structural biology. RSC Adv. 9, 8695–8699 (2019).

5. I. Schlichting, S. C. Almo, G. Rapp, K. Wilson, K. Petratos, A. Lentfer, A. Wittinghofer, W. Kabsch, E. F. Pai, G. A. Petsko, R. S. Goody, Time-resolved X-ray crystallographic study of the conformational change in Ha-Ras p21 protein on GTP hydrolysis. Nature 345, 309–315 (1990).

6. B. L. Stoddard, P. Koenigs, N. Porter, K. Petratos, G. A. Petsko, D. Ringe, Observation of the light-triggered binding of pyrone to chymotrypsin by Laue x-ray crystallography. Proceedings of the National Academy of Sciences 88, 5503–5507 (1991).

7. E. M. H. Duke, A. Hadfield, S. Walters, S. Wakatsuki, R. K. Bryan, L. N. Johnson, Time-resolved diffraction studies on glycogen phosphorylase b. Philos Trans A Math Phys Eng Sci 340, 245–261 (1992).

8. T. Tosha, T. Nomura, T. Nishida, N. Saeki, K. Okubayashi, R. Yamagiwa, M. Sugahara, T. Nakane, K. Yamashita, K. Hirata, G. Ueno, T. Kimura, T. Hisano, K. Muramoto, H. Sawai, H. Takeda, E. Mizohata, A. Yamashita, Y. Kanematsu, Y. Takano, E. Nango, R. Tanaka, O. Nureki, O. Shoji, Y. Ikemoto, H. Murakami, S. Owada, K. Tono, M. Yabashi, M. Yamamoto, H. Ago, S. Iwata, H. Sugimoto, Y. Shiro, M. Kubo, Capturing an initial intermediate during the P450nor enzymatic reaction using time-resolved XFEL crystallography and caged-substrate. Nat Commun 8, 1585 (2017).

9. P. Smyth, S. Jaho, L. J. Williams, G. Karras, A. Fitzpatrick, A. J. Thompson, S. Battah, D. Axford, S. Horrell, M. Lučić, K. Ishihara, M. Kataoka, H. Matsuura, K. Shimba, K. Tono, T. Tosha, H. Sugimoto, S. Owada, M. A. Hough, J. A. R. Worrall, R. L. Owen, Time-resolved serial synchrotron and serial femtosecond crystallography of heme proteins using photocaged nitric oxide. IUCrJ 12, 582–594 (2025).

10. P. Mehrabi, E. C. Schulz, R. Dsouza, H. M. Müller-Werkmeister, F. Tellkamp, R. J. D. Miller, E. F. Pai, Time-resolved crystallography reveals allosteric communication aligned with molecular breathing. Science 365, 1167–1170 (2019).

11. M. Wilamowski, D. A. Sherrell, Y. Kim, A. Lavens, R. W. Henning, K. Lazarski, A. Shigemoto, M. Endres, N. Maltseva, G. Babnigg, S. C. Burdette, V. Srajer, A. Joachimiak, Time-resolved β-lactam cleavage by L1 metallo-β-lactamase. Nat Commun 13, 7379 (2022).

12. M. Kikkawa, Y. Sasaki, S. Kawata, Y. Hatakeyama, F. B. Ueno, K. Saito, Photochemical and thermal decomposition of (ΔΔ,ΛΛ)-(μ-Hydroxo)(μ-peroxo)bis[bis(ethylenediamine)cobalt(III)] ions in basic aqueous solution. Inorg. Chem. 24, 4096–4100 (1985).

13. R. MacArthur, A. Sucheta, F. F. Chong, O. Einarsdóttir, Photodissociation of a (μ-peroxo)(μ-hydroxo)bis[bis(bipyridyl)-cobalt(III)] complex: a tool to study fast biological reactions involving O2. Proceedings of the National Academy of Sciences 92, 8105–8109 (1995).

14. Ó. Einarsdóttir, C. Funatogawa, T. Soulimane, I. Szundi, Kinetic studies of the reactions of O_2_ and NO with reduced *Thermus thermophilus ba*3 and bovine *aa*3 using photolabile carriers. Biochimica et Biophysica Acta (BBA) - Bioenergetics 1817, 672–679 (2012).

15. C. Ludovici, R. Fröhlich, K. Vogtt, B. Mamat, M. Lübben, Caged O_2_. European Journal of Biochemistry 269, 2630–2637 (2002).

16. A. R. Howard-Jones, V. Adam, A. Cowley, J. E. Baldwin, D. Bourgeois, Cryophotolysis of a caged oxygen compound for use in low temperature biological studies. Photochem Photobiol Sci 8, 1150–1156 (2009).

17. E. Sandelin, J. Johannesson, O. Wendt, G. Brändén, R. Neutze, C.-J. Wallentin, Characterization and evaluation of photolabile (µ-peroxo)(µ-hydroxo)bis[bis(bipyridyl)cobalt caged oxygen compounds to facilitate time-resolved crystallographic studies of cytochrome c oxidase. Photochem Photobiol Sci 23, 839–851 (2024).

18. J. Schulz, J. Bielecki, R. B. Doak, K. Dörner, R. Graceffa, R. L. Shoeman, M. Sikorski, P. Thute, D. Westphal, A. P. Mancuso, A versatile liquid-jet setup for the European XFEL. J Synchrotron Rad 26, 339–345 (2019).

19. A. M. Orville, N. Elango, J. D. Lipscomb, D. H. Ohlendorf, Structures of Competitive Inhibitor Complexes of Protocatechuate 3,4-Dioxygenase: Multiple Exogenous Ligand Binding Orientations within the Active Site,. Biochemistry 36, 10039–10051 (1997).

20. P. Rabe, J. H. Beale, A. Butryn, P. Aller, A. Dirr, P. A. Lang, D. N. Axford, S. B. Carr, T. M. Leissing, M. A. McDonough, B. Davy, A. Ebrahim, J. Orlans, S. L. S. Storm, A. M. Orville, C. J. Schofield, R. L. Owen, Anaerobic fixed-target serial crystallography. IUCrJ 7, 901–912 (2020).

21. M. Bjelčić, K. G. V. Sigfridsson Clauss, O. Aurelius, M. Milas, J. Nan, T. Ursby, Anaerobic fixed-target serial crystallography using sandwiched silicon nitride membranes. Acta Cryst D 79, 1018–1025 (2023).

22. A. Ostermann, R. Waschipky, F. G. Parak, G. U. Nienhaus, Ligand binding and conformational motions in myoglobin. Nature 404, 205–208 (2000).

23. V. Šrajer, T. Teng, T. Ursby, C. Pradervand, Z. Ren, S. Adachi, W. Schildkamp, D. Bourgeois, M. Wulff, K. Moffat, Photolysis of the Carbon Monoxide Complex of Myoglobin: Nanosecond Time-Resolved Crystallography. Science 274, 1726–1729 (1996).

24. T. R. M. Barends, A. Gorel, S. Bhattacharyya, G. Schirò, C. Bacellar, C. Cirelli, J.-P. Colletier, L. Foucar, M. L. Grünbein, E. Hartmann, M. Hilpert, J. M. Holton, P. J. M. Johnson, M. Kloos, G. Knopp, B. Marekha, K. Nass, G. Nass Kovacs, D. Ozerov, M. Stricker, M. Weik, R. B. Doak, R. L. Shoeman, C. J. Milne, M. Huix-Rotllant, M. Cammarata, I. Schlichting, Influence of pump laser fluence on ultrafast myoglobin structural dynamics. Nature 626, 905–911 (2024).

25. S. Jaho, D. Axford, D.-H. Gu, M. A. Hough, R. L. Owen, “Use of fixed targets for serial crystallography” in Methods in Enzymology (Academic Press, 2024), vol. 709, pp. 29–55.

26. R. L. Owen, D. Axford, D. A. Sherrell, A. Kuo, O. P. Ernst, E. C. Schulz, R. J. D. Miller, H. M. Mueller-Werkmeister, Low-dose fixed-target serial synchrotron crystallography. Acta Cryst D 73, 373–378 (2017).

27. G. Gotthard, A. Flores-Ibarra, M. Carrillo, M. W. Kepa, T. J. Mason, D. P. Stegmann, B. Olasz, M. Pachota, F. Dworkowski, D. Ozerov, B. F. Pedrini, C. Padeste, J. H. Beale, P. Nogly, Fixed-target pump–probe SFX: eliminating the scourge of light contamination. IUCrJ 11, 749–761 (2024).

28. J. J. A. G. Kamps, P. Hinchliffe, J. Glerup, E. I. Freeman, P. A. Lang, C. L. Tooke, M. Beer, L. Parkinson, D.-H. Gu, S. Park, N. Devenish, T. Zhou, A. Shilova, S. Kaur, P. Rabe, C. J. Schofield, J. Spencer, J. Park, R. L. Owen, A. M. Orville, P. Aller, Drop-on-fixed-target reaction initiation approach for serial and time-resolved crystallography. IUCrJ 13, 471–484 (2026).

29. J. L. Dickerson, P. T. N. McCubbin, J. C. Brooks-Bartlett, E. F. Garman, Doses for X-ray and electron diffraction: New features in RADDOSE-3D including intensity decay models. Protein Science 33, e5005 (2024).

30. J. L. Dickerson, P. T. N. McCubbin, E. F. Garman, RADDOSE-XFEL: femtosecond time-resolved dose estimates for macromolecular X-ray free-electron laser experiments. J Appl Cryst 53, 549–560 (2020).

31. L. J. Williams, A. J. Thompson, P. Dijkstal, M. Appleby, G. Assmann, F. S. N. Dworkowski, N. Hiller, C.-Y. Huang, T. Mason, S. Perrett, E. Prat, D. Voulot, B. Pedrini, J. H. Beale, M. A. Hough, J. A. R. Worrall, R. L. Owen, Damage before destruction? X-ray-induced changes in single-pulse serial femtosecond crystallography. IUCrJ 12, 358–371 (2025).

32. T. Nakane, Y. Joti, K. Tono, M. Yabashi, E. Nango, S. Iwata, R. Ishitani, O. Nureki, Data processing pipeline for serial femtosecond crystallography at SACLA. J Appl Cryst 49, 1035–1041 (2016).

33. J. Beilsten-Edmands, J. M. Parkhurst, G. Winter, G. Evans, “Processing serial synchrotron crystallography diffraction data with DIALS” in Methods in Enzymology, (Academic Press, 2024), vol. 709, pp. 207–244.

34. T. A. White, R. A. Kirian, A. V. Martin, A. Aquila, K. Nass, A. Barty, H. N. Chapman, CrystFEL: a software suite for snapshot serial crystallography. J Appl Cryst 45, 335–341 (2012).

35. A. Barty, R. A. Kirian, F. R. N. C. Maia, M. Hantke, C. H. Yoon, T. A. White, H. Chapman, Cheetah: software for high-throughput reduction and analysis of serial femtosecond X-ray diffraction data. J Appl Cryst 47, 1118–1131 (2014).

36. Y. Gevorkov, O. Yefanov, A. Barty, T. A. White, V. Mariani, W. Brehm, A. Tolstikova, R.-R. Grigat, H. N. Chapman, XGANDALF – extended gradient descent algorithm for lattice finding. Acta Cryst A 75, 694–704 (2019).

37. T. A. White, Detector alignment for X-ray crystallography using Millepede-II. J Appl Cryst 59, 594–608 (2026).

38. T. A. White, V. Mariani, W. Brehm, O. Yefanov, A. Barty, K. R. Beyerlein, F. Chervinskii, L. Galli, C. Gati, T. Nakane, A. Tolstikova, K. Yamashita, C. H. Yoon, K. Diederichs, H. N. Chapman, Recent developments in CrystFEL. J Appl Cryst 49, 680–689 (2016).

39. J. Beilsten-Edmands, G. Winter, R. Gildea, J. Parkhurst, D. Waterman, G. Evans, Scaling diffraction data in the DIALS software package: algorithms and new approaches for multi-crystal scaling. Acta Cryst D 76, 385–399 (2020).

40. A. J. McCoy, R. W. Grosse-Kunstleve, P. D. Adams, M. D. Winn, L. C. Storoni, R. J. Read, Phaser crystallographic software. J Appl Cryst 40, 658–674 (2007).

41. A. Vagin, A. Teplyakov, Molecular replacement with MOLREP. Acta Cryst D 66, 22–25 (2010).

42. T. R. M. Barends, L. Foucar, A. Ardevol, K. Nass, A. Aquila, S. Botha, R. B. Doak, K. Falahati, E. Hartmann, M. Hilpert, M. Heinz, M. C. Hoffmann, J. Köfinger, J. E. Koglin, G. Kovacsova, M. Liang, D. Milathianaki, H. T. Lemke, J. Reinstein, C. M. Roome, R. L. Shoeman, G. J. Williams, I. Burghardt, G. Hummer, S. Boutet, I. Schlichting, Direct observation of ultrafast collective motions in CO myoglobin upon ligand dissociation. Science 350, 445–450 (2015).

43. L. Potterton, J. Agirre, C. Ballard, K. Cowtan, E. Dodson, P. R. Evans, H. T. Jenkins, R. Keegan, E. Krissinel, K. Stevenson, A. Lebedev, S. J. McNicholas, R. A. Nicholls, M. Noble, N. S. Pannu, C. Roth, G. Sheldrick, P. Skubak, J. Turkenburg, V. Uski, F. von Delft, D. Waterman, K. Wilson, M. Winn, M. Wojdyr, CCP4i2: the new graphical user interface to the CCP4 program suite. Acta Cryst D 74, 68–84 (2018).

44. G. N. Murshudov, P. Skubák, A. A. Lebedev, N. S. Pannu, R. A. Steiner, R. A. Nicholls, M. D. Winn, F. Long, A. A. Vagin, REFMAC5 for the refinement of macromolecular crystal structures. Acta Cryst D 67, 355–367 (2011).

45. P. D. Adams, P. V. Afonine, G. Bunkóczi, V. B. Chen, I. W. Davis, N. Echols, J. J. Headd, L.-W. Hung, G. J. Kapral, R. W. Grosse-Kunstleve, A. J. McCoy, N. W. Moriarty, R. Oeffner, R. J. Read, D. C. Richardson, J. S. Richardson, T. C. Terwilliger, P. H. Zwart, PHENIX: a comprehensive Python-based system for macromolecular structure solution. Acta Cryst D 66, 213–221 (2010).

46. P. Emsley, B. Lohkamp, W. G. Scott, K. Cowtan, Features and development of Coot. Acta Cryst D 66, 486–501 (2010).

47. D. Liebschner, P. V. Afonine, N. W. Moriarty, B. K. Poon, O. V. Sobolev, T. C. Terwilliger, P. D. Adams, Polder maps: improving OMIT maps by excluding bulk solvent. Acta Cryst D 73, 148–157 (2017).

48. V. B. Chen, W. B. Arendall, J. J. Headd, D. A. Keedy, R. M. Immormino, G. J. Kapral, L. W. Murray, J. S. Richardson, D. C. Richardson, MolProbity: all-atom structure validation for macromolecular crystallography. Acta Cryst D 66, 12–21 (2010).

49. R. P. Joosten, F. Long, G. N. Murshudov, A. Perrakis, The PDB_REDO server for macromolecular structure model optimization. IUCrJ 1, 213–220 (2014).

50. J. Y. Young, J. D. Westbrook, Z. Feng, R. Sala, E. Peisach, T. J. Oldfield, S. Sen, A. Gutmanas, D. R. Armstrong, J. M. Berrisford, L. Chen, M. Chen, L. Di Costanzo, D. Dimitropoulos, G. Gao, S. Ghosh, S. Gore, V. Guranovic, P. M. S. Hendrickx, B. P. Hudson, R. Igarashi, Y. Ikegawa, N. Kobayashi, C. L. Lawson, Y. Liang, S. Mading, L. Mak, M. S. Mir, A. Mukhopadhyay, A. Patwardhan, I. Persikova, L. Rinaldi, E. Sanz-Garcia, M. R. Sekharan, C. Shao, G. J. Swaminathan, L. Tan, E. L. Ulrich, G. van Ginkel, R. Yamashita, H. Yang, M. A. Zhuravleva, M. Quesada, G. J. Kleywegt, H. M. Berman, J. L. Markley, H. Nakamura, S. Velankar, S. K. Burley, OneDep: Unified wwPDB System for Deposition, Biocuration, and Validation of Macromolecular Structures in the PDB Archive. Structure 25, 536–545 (2017).

51. H. Yang, V. Guranovic, S. Dutta, Z. Feng, H. M. Berman, J. D. Westbrook, Automated and accurate deposition of structures solved by X-ray diffraction to the Protein Data Bank. Acta Cryst D 60, 1833–1839 (2004).

52. D.-H. Gu, D. Axford, J. Beilsten-Edmands, S. Jaho, R. L. Owen, Increasing X-ray energy improves data quality in serial crystallography. J Synchrotron Rad 33, 344–350 (2026).

53. D. von Stetten, A. R. Pearson, Decision-making in serial crystallography: a simple test to quickly determine whether sufficient data have been collected. Acta Cryst D 82, 227–236 (2026).

